# Mapping Bidirectional Allosteric Communication in Arf GTPases

**DOI:** 10.1101/2025.11.21.689788

**Authors:** Edgar Peters, Tejaswi Koduru, Scott A. McCallum, Estella F. Yee, Jacqueline Cherfils, Catherine A. Royer

## Abstract

The Arf GTPases are eukaryotic signaling proteins implicated in trafficking, motility and membrane remodeling. They must undergo a massive conformational transition in the switch between their inactive, GDP-bound form and their GTP-bound form, competent for downstream signaling. The mechanism of their GDP-to-GTP nucleotide switch implicates a functional molten globule (MG) ensemble. Access to this ensemble and apparent spontaneous switching is modulated by residues in the C-terminal half of the different homologs through a back-to-front allosteric pathway. Here, using high pressure (HP) NMR we show that a mutation in the N-terminal switch region in the front of Arf1 known to modulate spontaneous switching perturbs the stability of residues on a path that stretches from the front to the back. This establishes the existence of a bidirectional and continuous allosteric pathway that passes through the GDP ligand. HP switching studies also demonstrate that the allosteric mechanism controls access to the functional MG state, rather than directly affecting the switching rates.

**Secondary Abstract:** The mechanism of the Arf GTPase nucleotide switch implicates a functional molten globule (MG) ensemble. Switching by the front side of the Arf proteins is controlled by sequences on the back side. Here, using high pressure (HP) NMR we map a bidirectional and continuous allosteric pathway that passes between front and back through the GDP ligand. We also demonstrate that the allosteric mechanism controls access to the functional MG state, rather than directly affecting the switching rates.

## Introduction

ADP ribosylation factors (Arfs) are small GTPases that constitute a subfamily within the larger family of Ras GTPases (1, 2). Highly conserved from yeast to humans (3–5), Arfs are involved in vesicle trafficking and membrane remodeling, playing important roles in motility and signaling (6–8), and thus in human disease (7, 9). Like all small GTPases, they switch conformations between a GDP-bound state which does not lead to downstream signaling and a GTP-bound state that is signaling-competent. Signaling is turned off by the hydrolysis of GTP to GDP, which returns the Arfs to the off state. Both events are intrinsically extremely slow or even totally inhibited, and hence require regulatory proteins (1, 10). To efficiently release GDP and bind GTP, which is present in cells at much higher concentrations than GDP, Arfs must interact with Guanosine nucleotide Exchange Factors (GEFs). Likewise, efficient hydrolysis requires interaction with a GTPase Activating Protein (GAP) (1, 10).

Arfs distinguish themselves from the other subfamilies of small GTPases by the very large conformational change between the GDP– and GTP-bound forms (Figure S1) (3, 11–14). Upon switching from the dinucleotide to the trinucleotide bound forms, the N-terminal helix dissociates from the core of the protein. This N-terminal helix is myristoylated *in vivo* and interacts with membranes when it is released from the core (15, 16), thus localizing Arf functions to membranes. The switch 1 region peels off from the central β-sheet and moves several Å to interact with the GTP ligand. The interswitch β-hairpin undergoes a 2-residue shift in register into the space previously occupied by the N-terminal helix in the GDP-bound state. Finally, the switch 2 region becomes more ordered and moves to interact with the γ-phosphate of the GTP ligand.

Arf1 and Arf6 are the most well-known members of the Arf and Arf-like (Arl) subfamily. They are highly homologous with over 95% sequence similarity and 70% sequence identity (Figure S2). Arf1 functions mostly at the Golgi membranes (17, 18) whereas Arf6 is found at the plasma membrane functioning in trafficking and cytoskeletal and signaling pathways (6, 8, 19–21). In addition to distinct cellular localization, Arf1 and Arf6 interact with different signaling partners and small molecules (22–25). Arf1 does not exhibit significant spontaneous switching even at very low Mg^2+^ concentrations, whereas Arf6 can undergo the GDP to GTP switch at 1 μM Mg^2+^, albeit slowly (26).

Using NMR and other biophysical techniques coupled to high hydrostatic pressure (HP), we recently demonstrated that the mechanism for the nucleotide switch transition in both Arf1 and Arf6 implicates a functional molten globule (MG) ensemble, which is less stable than the native GDP-bound state (26–28). Accessing this functional MG primes the Arfs to undergo the highly disruptive and energetically costly conformational changes implicated in the switch transition itself. Our recent work also revealed that residues in helix 5 on the backside, where many of the sequence differences between Arf1 and Arf6 are located (Figure S2), modulate the spontaneous switching rate of the Arfs (26, 27), revealing a back-to-front control pathway. While this recent work yielded important insights into the mechanisms and sequence determinants of the nucleotide switch transition in the Arf proteins (26–28), the pathways of allosteric communication and their differences among Arf homologs remain to be fully described.

Following thermodynamic principles, the recently elucidated Arf back-to-front allosteric pathway (26) must be bi-directional. However, how residues in the switch region are implicated in reverse front-to-back allosteric communication and how they energetically impact the Arf conformational landscape is not known. To address these questions, we turned to a previously studied substitution between Arf1 (residue I42) and Arf6 (equivalent residue S38) (13). This residue, which is located on the front of Arf-GDP structures, was shown to be an important determinant of the GDP/GTP switch rate. The crystal structures of Arf1-GDP (3) and Arf6-GDP (13) revealed that the I/S substitution leads to significant differences near the expected Mg^2+^ ion (13) (Figure 1A, B). In Arf1-GDP, residue I42 interacts with L34 in the facing helix 2, while nearby E54 in the interswitch coordinates the Mg^2+^ ion, along with the invariant T31 near the P-loop (Figure 1A). The positioning of residue E54 is further stabilized by packing against V65. In contrast, in Arf6-GDP the hydroxyl group of the S38 sidechain attracts the E50 side chain outside the Mg^2+^ coordination sphere to form an alternative hydrogen bond (Figure 1B), and Mg^2+^ was not observed in the crystal structure where it was modeled instead by an NH4^+^ ion. The I42S mutation in Arf1 deleted for the N-terminal helix, Arf1Δ17, increased the spontaneous switching rate to that of the corresponding deletion mutant of Arf6, Arf6Δ13. Conversely, the S38I mutation in Arf6Δ13 led to slower switching, at a rate equivalent to that of Arf1Δ17 (13). Thus, the I/S substitution in Arf6 was proposed to decrease the Mg^2+^-GDP affinity and result in faster switching (13). We therefore reasoned that the I42S Arf1 mutant should be well-suited to study the reverse allosteric communication in Arf1.

**Figure 1.**
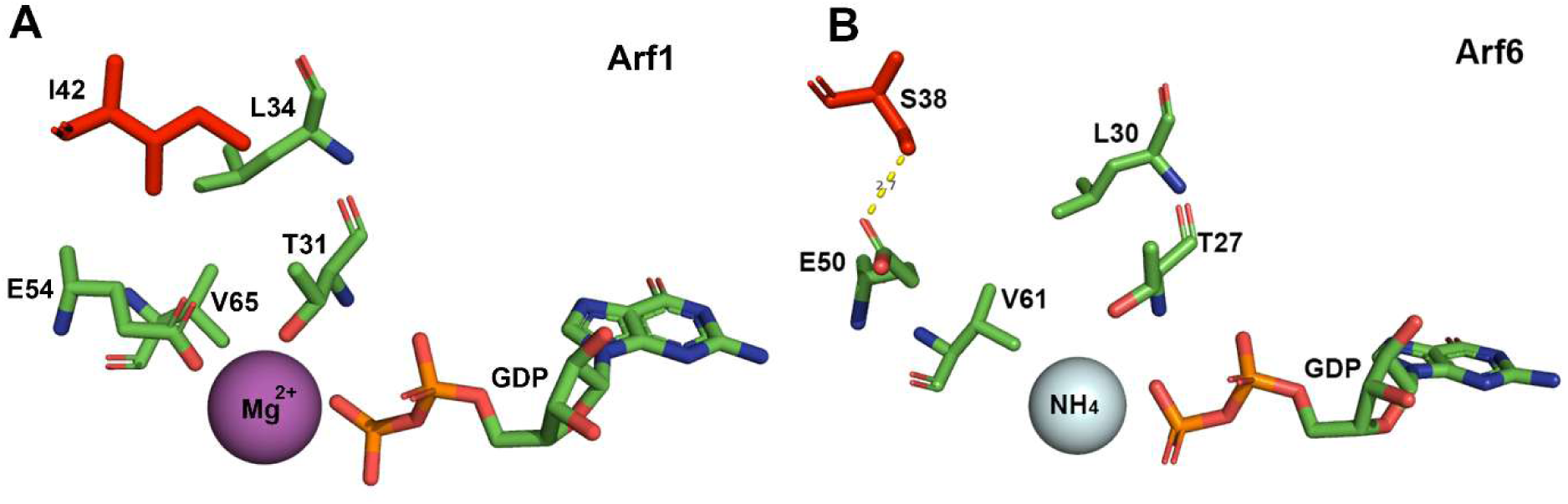
The interaction network implicating I42 in Arf1 is disrupted in Arf6. A) The interaction network of I42 in Arf1. I42 interacts with L34 in the opposing helix 2. This allows E54 to coordinate the Mg^2+^ ion, along with T31. E534 also packs against V65. B) The interaction network of S38 in Arf6. S38 in Arf6 is equivalent to I42 in Arf1, given that helix 1 in Arf6 is shorter by 4 residues than in Arf1. E50 in Arf6 (equivalent to E54 in Arf1) does not coordinate the Mg^2+^ ion. Rather it forms an H-bond with S38. interacts with L34 in the opposing helix 2. The specified residues are labeled and shown in CPK stick representation. The GDP ligand is also shown in CPK stick representation. The Mg^2+^ ion is magenta.

In this study, we used we used HP NMR coupled with other biophysical approaches (26–28) to establish the impact of the I42S mutation in Arf1 on the residue-specific stability distribution. We found that the I42S mutation in Arf1 significantly perturbed local stability along a pathway of communication from the switch region to the P-loop, through the GDP ligand, its interacting loops, and finally to the backside of the protein in helices 4 and 5. The perturbation of residues on the backside of the protein by mutations in the front demonstrates that the previously identified back-to-front allosteric pathway (26) is bidirectional, as required by thermodynamics. Moreover, the pressure dependence of switching rates of the WT and I42S Arf1 revealed that the mechanism of the mutation in increasing switching probability is to favor the population of the functional molten globule, rather than impacting the switch transition itself.

## Material and Methods

### Protein production and purification

Full-length Arf1 and Arf1 I42S were produced in *E. coli* BL21 (DE3) cells in LB or minimal media as an N-terminal His-Tag fusion bearing a thrombin cleavage site using the pET-15b plasmid. After disrupting the cells and collecting the supernatant, the solution was passed over a Ni-NTA column. After loading, the column was washed in 20 mM Bis-Tris, 1 mM TCEP, 10mM NaCl, 10mM imidazole, pH 7.4. Then Arf1 variants were eluted using the same buffer with 250 mM imidazole. The His-Tag was removed by thrombin cleavage at 22 °C for 18-20 hours. 1 mM PMSF was added to stop the reaction. Cleavage was verified by SDS PAGE and Maldi mass spectrometry. After thrombin cleavage of the His-Tag, only two amino acids (Gly-Ser) remain N-terminal to Arf1 residue 1. Arf1-GDP was further purified by size exclusion chromatography on an S75 Sepharose column.

Variants of WT Arf1 and I42S in which the N-terminal helix was deleted, Arf1Δ17 and Arf1Δ17 I42S, were produced as C-terminal His-Tag fusions from a pET 24a+ vector with an additional arginine residue C-terminal to the Arf1 sequence. After purification on the Ni-NTA column, as above, the C-terminal His-Tag was cleaved with carboxypeptidase A (Sigma-Aldrich) in 50 mM Tris, 1 mM MgCl2, and 150 mM NaCl overnight at pH 7.5. Cleavage was verified by SDS PAGE and Maldi mass spectrometry. Cleaved Arf1Δ17 or Arf1Δ17 I42S was buffer exchanged to 50 mM bis-Tris, 150 mM NaCl, 1 mM MgCl2, and 5 mM DTT at pH 6.5. Then GDP exchange was performed on Arf1Δ17, Arf1 I42S and Arf1Δ17 I42S (29) to obtain 100% GDP-bound protein samples. The final step was gel filtration using a Hi-Load 16/600 S75 Sepharose column.

### NMR backbone amide assignments

Amide backbone resonance assignments for WT Arf1 were previously determined (27) and the established chemical shift data in addition to correlations detected in 2D 1H-15N HSQC, and 3D ^15^N NOESY– and TOCSY-HSQC NMR experiments performed on WT Arf1 and the Arf1 I42S mutant constructs were used to assign I42S Arf1 amide resonances Spectra were acquired on samples containing 0.5 mM WT Arf1 or Arf1 I42S in 50 mM bis-Tris, 150 mM NaCl, 1 mM MgCl2, and 5 mM DTT at pH 6.5 at 20◦C on a 600 MHz Bruker NEO NMR spectrometer. A comparison of the WT and I42S mutant HSQC spectra is shown in Figure S3, along with an example of the NOESY and TOCSY spectra used for resonance assignments of residues with large chemical shift perturbations in the mutant. The amide resonances for a total of 134 residues were resolved and confidently assigned in the I42S 1H-15N HSQC spectrum (Table S1).

### High pressure NMR data acquisition and analysis

HP NMR was carried out as previously described (27) using the ceramic tube developed by Peterson and Wand (30) (Daedalus Innovations, Aston, PA). HP NMR was carried out in the same buffer as above as previously described (27). For the I42S mutant, of the peaks that could be quantified as a function of pressure, 112 had corresponding *ΔG_app_* values available in the Arf1 HP NMR data sets at pH 6.5 (27) for comparison.

For the analysis of the pressure-induced intensity loss transitions, for any given pressure, each residue in a protein ensemble is either in its native state environment or it is not, depending on the change in stability and volume between the native and non-native like conformations. Thus, we fit profiles for the pressure-induced reduction in native state backbone amide ^1^H-^15^N HSQC peak intensities to a two-state model as follows:

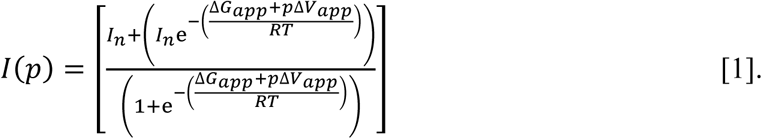

Here *I_p_* and *I_n_* represent the intensity at pressure, *p*, and the intensity of the native state at atmospheric pressure, respectively. For this analysis, if necessary, the low-pressure plateau value was fixed as 105% of the average of the first two pressure points, taken at atmospheric pressure and 10 bar. For a few residues with no apparent low pressure plateau, the value was constrained to the average of the atmospheric pressure intensity for all residues. We constrained the high-pressure plateau, corresponding to 100% population of the excited state, to 0, representing a full depopulation of the native state ensemble and thus no native state peak intensity. The apparent change in the free energies (*ΔG_app_*) and the molar volume (*ΔV_app_*) for the transition from native to excited state ensembles was determined on a residue-specific basis by least-squares fitting to the above equation, done with a suite of in-house MATLAB scripts.

### High pressure SAXS

HP-SAXS experiments were conducted on GDP-loaded Arf1Δ17 at the ID7A beamline at the Cornell High Energy Synchrotron source, CHESS, at pH 6.5 as previously described (27). The 2D SAXS profiles were processed using BioXTAS RAW 2.1.4 package (31). The radius of gyration was determined using Guinier analysis and the pair distance distribution function (P(r)), computed with the GNOM package from the ATSAS 3.0.04-2 suite, integrated into the RAW software (32).

### High-pressure fluorescence measurement of nucleotide exchange

Time-dependent HP fluorescence was carried out in a home-built HP optical cell using a modified ISS Koala fluorometer (ISS, Inc., Champaign, IL) as previously described (34), and an automated pump (Pressure Biosciences, Canton, MA). The buffer was the same as for the HP NMR experiments, and Arf1Δ17 was at a concentration of 50 μM. The exchange of GDP by GTP was measured using 20 μM of GDP-loaded Arf1Δ17 at pH 6.5 with 50 mM excess GTP at 33◦C as previously described (13). A 120-s dead-time was required before data acquisition for sample loading and pressurization

## Results

### The Arf1 I42S mutant populates a broad MG ensemble under pressure

We first sought to establish the effects of the I42S mutation on Arf1 residue-specific stability. To do so, we turned to HP NMR of Arf1 I42S-GDP in the presence of 1 mM Mg^2+^ to circumvent its low affinity. Due to significant chemical shift perturbations of several amide crosspeaks resulting from the mutation, the ^1^H-^15^N HSQC spectrum of the mutant was assigned based on ^15^N NOESY and TOCSY experiments (Table S1, Figure S3). Next, NMR ^1^H-^15^N HSQC spectra of the Arf1 I42S variant were acquired as a function of pressure. The intensities of the native state backbone amide crosspeaks exhibited pressure-dependent decreases in intensity over a broad pressure range, with no appearance of unfolded state peaks (Figure 2A). In addition, residues in the switch region were more labile under pressure than those in the C-terminal half of the protein Residue-specific apparent stabilities (*ΔG_app_*) and volume changes (*ΔV_app_*) were obtained from fits of the transition curves to a 2-state transition for each residue that could be quantified as a function of pressure (Table S2, Figure 2B, C, Figure S4). The loss of native state peak intensity over a broad pressure range with no concomitant increase in unfolded state peaks was shown previously for both Arf1 and Arf6 to be due to the population of a broad MG ensemble undergoing conformational exchange on the NMR timescale (26, 27). Thus, the present HP NMR results indicate that Arf1 I42S-GDP displays an MG ensemble at high pressure, whose properties can be analyzed in comparison with those of WT Arf1.

**Figure 2.**
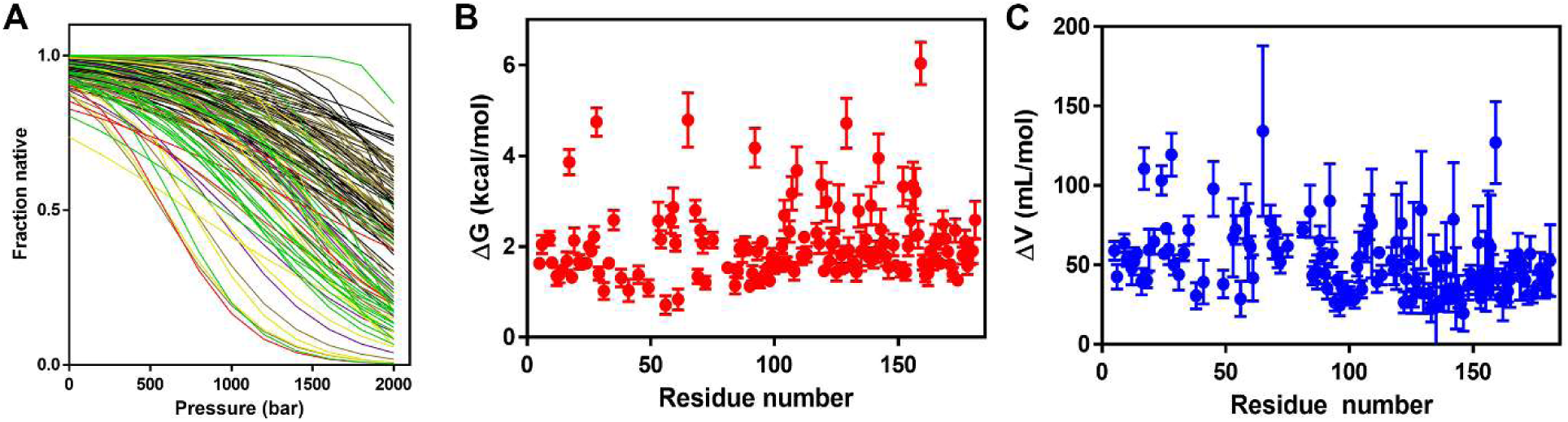
Arf1 I42S forms a molten globule excited state at high pressure. A) Fits of the pressure dependent decrease in intensity of the ^1^H-^15^N native state amide HSQC peaks. Data and fits can be found in supplemental File 1. B) The residue-specific stabilities, ΔG_app_ and C) ΔV_app_, respectively for Arf1 I42S recovered from the fits shown in A.

To further confirm that Arf1 I42S-GDP remains globular at high pressure, we used small angle X-ray scattering (SAXS), which provides information about the protein size and shape. HP SAXS demonstrated that despite losing most of the native state amide NMR crosspeak intensity, Arf1 I42S remained mostly globular at high pressure (Figure 3, Figure S5), as was found previously for both WT Arf1 and Arf6 (26, 27). Comparison of the normalized Kratky plots of the scattering intensities show that the I42S mutant has a somewhat larger size than WT Arf1 at atmospheric pressure (Figure 3A). This increase in size was slightly larger at 3000 bar (Figure 3B), although Arf1 I42S remained globular. This can be explained by the fact that while the I42S mutant expanded slightly at this pressure (Figure 3C), WT Arf1 contracted slightly (Figure 3D). Overall, the HP NMR and HP SAXS results are consistent with the population of an MG ensemble for the I42S mutant of Arf1.

**Figure 3.**
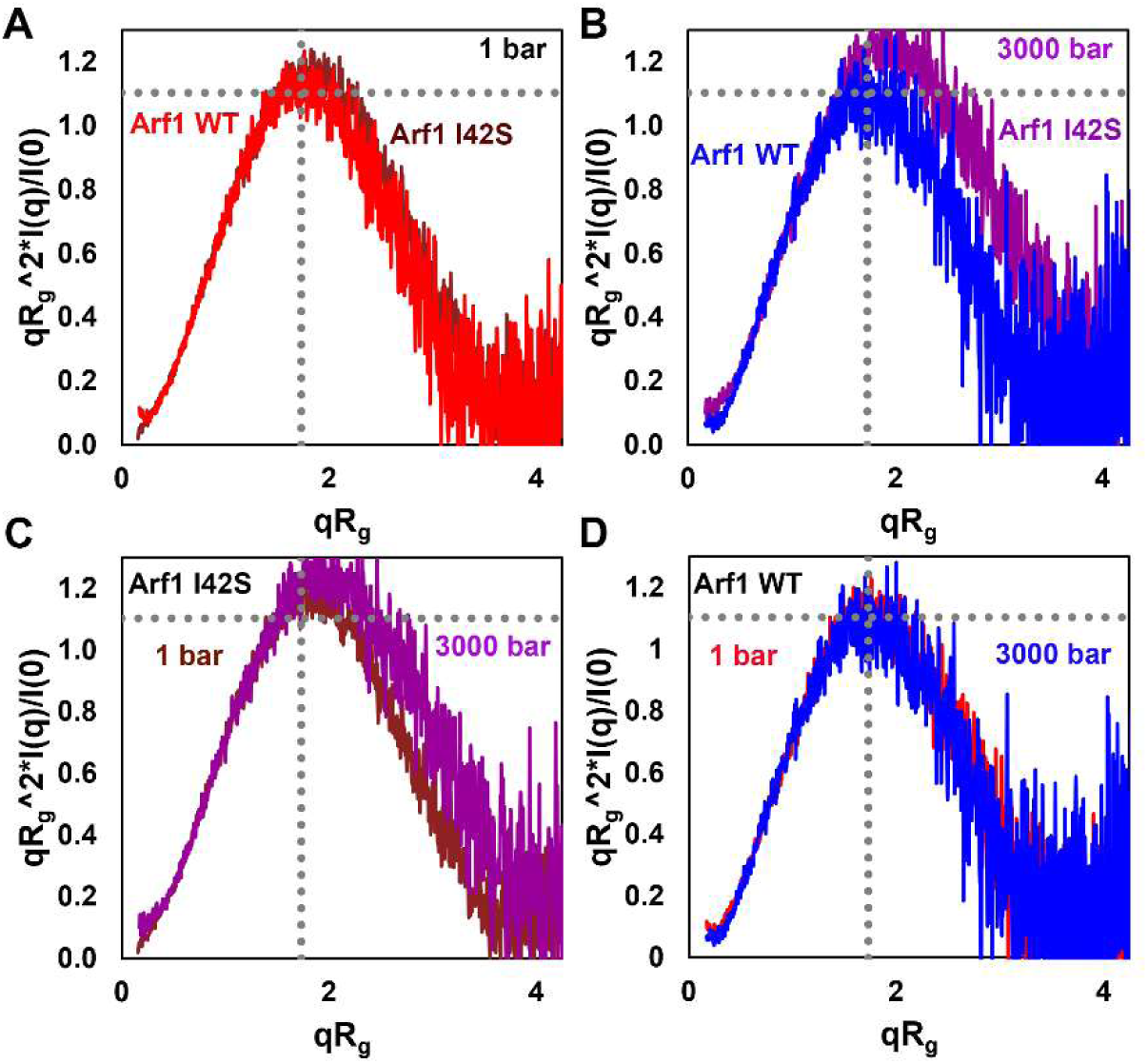
The shape of Arf1 I42S differs from that of WT Arf1. A) Dimensionless Kratky plot constructed from the SAXS intensity profiles for WT Arf1 (red) and Arf1 I42S (brown) at atmospheric pressure. B) Dimensionless Kratky plot constructed from the SAXS intensity profiles for WT Arf1 (red) and Arf1 I42S (brown) at 1500 bar. C) Dimensionless Kratky plot constructed from the SAXS intensity profiles for (red) Arf1 I42S at atmospheric pressure (brown) and at 3000 bar (purple). D) Dimensionless Kratky plot constructed from the SAXS intensity profiles for WT Arf1 at atmospheric pressure (red) and at 3000 bar (blue).

### The I42S mutation facilitates access to the molten globule, rather than enhancing the switching rate itself

We next sought to establish the effects of pressure on the switching kinetics of Arf1 I42S. Since Arf1Δ17, which lacks the first 17 residues, undergoes measurable spontaneous switching (16) while WT Arf1 does not, we used the Δ17 variants of WT Arf1 and Arf1 I42S to compare their nucleotide exchange kinetics by monitoring their intrinsic tryptophan fluorescence. As previously reported (13), we found that at atmospheric pressure Arf1Δ17 I42S switched faster than WT Arf1Δ17 (Figure 4A). Increasing the pressure to 1500 bar resulted in a faster switch for the I42S mutant (Figure 4B). This indicates that the MG state populated under pressure by Arf1Δ17 I42S, as in the case of WT Arf1Δ17 (27) and Arf6 (26), is functionally implicated in the nucleotide switch. Note that pressure also enhanced switching in WT Arf1Δ17 (Figure 4C), as shown previously (27, 28).

**Figure 4.**
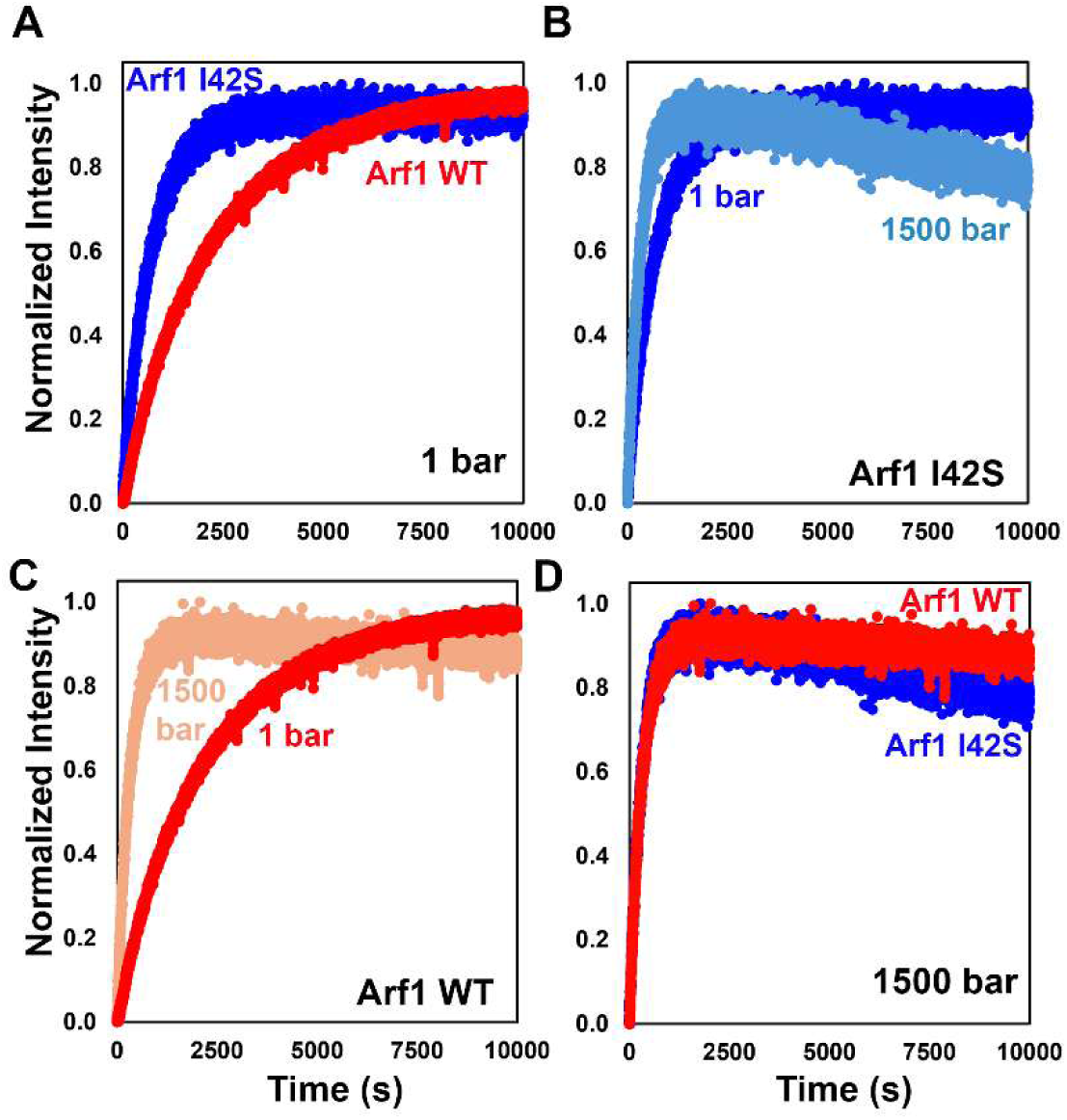
Pressure leads to identical switching for Arf1Δ17I42S and WT Arf1Δ17. A) Comparison of the fluorescence detected progress curves for the GDP/GTP switch for Arf1Δ17 I42S (blue) and WT Arf1Δ17 (red) at atmospheric pressure. B) Comparison of the fluorescence detected progress curves for the GDP/GTP switch for Arf1Δ17 I42S at atmospheric pressure (blue) and at 1500 bar (light blue). C) Comparison of the fluorescence detected progress curves for the GDP/GTP switch for WT Arf1Δ17 at atmospheric pressure (red) and at 1500 bar (light red). D) Comparison of the fluorescence detected progress curves for the GDP/GTP switch for Arf1Δ17 I42S (blue) and WT Arf1Δ17 (red) at 1500 bar.

Surprisingly, the switching rates for the WT Arf1Δ17 and Arf1Δ17 I42S were equivalent at high pressure (Figure 4D), in striking contrast with atmospheric pressure (compare with Figure 4A). We proposed previously that the MG excited state ensemble facilitates access to the higher energy conformations required for the GDP/GTP switch (27). The fact that switching occurs at the same rate for the mutant and WT suggests that the MG state is equally populated for both proteins at 1500 bar. Note as well that the GTPase activity of Arf1Δ17 I42S (decrease in fluorescence intensity at longer times) is enhanced under pressure compared to that of WT Arf1Δ17, which shows almost no GTPase activity even at high pressure (Figure 4B). The mechanism for the pressure-induced enhancement of spontaneous GTPase activity will be investigated elsewhere.

### The I42S mutation defines a front to back pathway

The I42S mutant of Arf1 populates the functional MG state and exhibits modified exchange kinetics. Hence, we sought to establish the impact of the I42S mutation on energetic communication within the Arf protein, including on the previously identified back-to-front pathway (26). To do so, we compared the residue specific stabilities extracted from fits of the NMR profiles of Arf1 I42S (Table S2, Figure S4) with those of WT Arf1 (27). This comparison revealed that 17 residues exhibited significant differences in their local stability (ΔΔG_app_ > ±1.5 kcal/mol) with respect to WT Arf1 (Figure 5A, Table 1, Table S3), two-thirds of which were destabilized in the I42S variant relative to WT Arf1.

**Figure 5.**
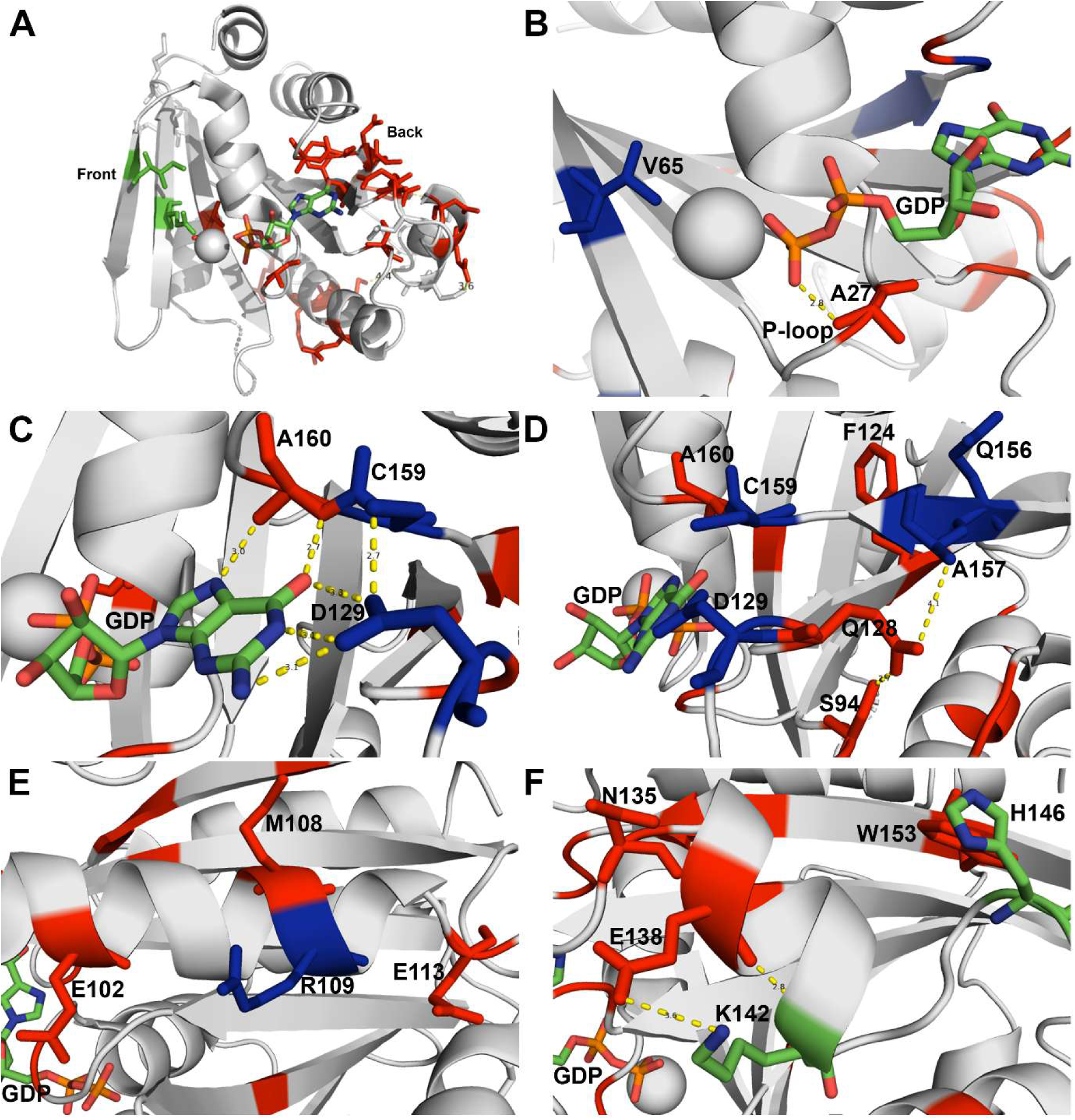
A bi-directional allosteric pathway links the switch region and the back side of Arf GTPases. A) Overall view of the Arf1 residues for which the local stability was perturbed by the I42S mutation. All perturbed residues are shown as red sticks regardless of the sign (positive or negative) of the energetic perturbation. The Mg^2+^ ion is in grey sphere, as is the cartoon of the rest of the protein. GDP is in CPK sticks. I42S (the location of the mutation) and E54 are shown in green sticks. The green arrow indicates the allosteric pathway. B-F) Zooms of the regions encompassing the perturbed residues. Stabilized residues are shown in blue sticks while destabilized residues are shown in red sticks. Yellow dashes indicate H-bonding interactions. Residue identities are as indicated. B) switch region and P-loop on the front side of the protein, C) The guanine-binding loops, D) Regions abutting the guanine-binding loops, E) Helix 4, F) Helix 5. Shown also is the destabilized W153 which interacts with H146 just after the kink in helix 5.

**Table 1.** Residues with significantly perturbed stability in Arf1I42S compared to Arf1 WT.

|  |  |  |
| --- | --- | --- |
| **Less stable in I42S** | **$\Delta\Delta G$ (kcal/mol)** |  |
| A27 | -1.6 | In the P-loop coordinating the β-phosphate in GDP |
| S94 | -1.8 | In the loop preceding helix H4 that packs against the P-loop, interacts with Q 128 |
| E102 | -2.5 | In helix H4 |
| M108 | -3.1 | In helix H4 |
| E113 | -1.8 | C-terminal to helix H4 |
| F124 | -3.1 | Center of central β-strand 6 |
| Q128 | -1.7 | At the end of β-strand 6, right before loop that contacts GDP |
| N135 | -2.1 | At the beginning of helix H5 |
| E138 | -1.6 | In the first turn of helix H5 |
| W153 | -2.8 | Tryptophan that packs against the leucine at the kink in helix H5 |
| A160 | -1.9 | Backbone NH H-bonds to the hydroxyl at position 6 in the guanine base |
| **More stable in I42S** |  |  |
| V65 | 2.8 | Packs against the E54 that coordinates $Mg^{2+}$ and H-bonds to T31 in WT Arf1-GDP. Packing is shifted away from M18 in Arf6 |
| R109 | 2.3 | Helix H4, exposed on surface |
| D129 | 3.4 | Coordinates N1 of guanosine base, may better coordinate given loss in A160 |
| Q156 | 1.6 | In β-strand 7, likely compensatory rearrangements due to destabilization of residues 126 and 128 |
| A157 | 1.5 | In β-strand 7, likely compensatory rearrangements due to destabilization of residues 126 and 128 |
| C159 | 4.4 | Just after β-strand 7, likely compensatory rearrangements due to destabilization of A160 |

Of all the perturbed residues, only A27 and V65 are located in the vicinity of the mutation (Figure 5B). A27 belongs to the conserved phosphate-binding P-loop of small GTPases and is close to the GDP phosphates. Its local stability decreased in the I42S variant relative to the WT protein. Assuming that the I42S mutant rearranges structurally to resemble Arf6 (Figure 1), this would remove the side chain of E54 from coordinating the Mg^2+^ ion, presumably destabilizing the Mg^2+^, and by association, the β-phosphate and A27. V65 is located in the interswitch, where it wedges between M22 and E54. Repositioning E54 to hydrogen bond with the S42 mutation, as in Arf6 (S38, Figure 1B), would disrupt its interactions with V65. Why this rearrangement results in stabilization of V65 may be entropic in origin. Of note, V65 was one of the residues that exhibited a large chemical shift perturbation in the ^1^H-^15^N HSQC spectrum with respect to WT (Figure S3).

Interestingly, three of the energetically perturbed residues (D129, C159, and A160) interact directly or indirectly with the guanine base, at the other end of the GDP nucleotide (Figure 5C). All three form hydrogen bonds with the guanine base, but only A160 was destabilized. This modulation of the GDP base interactions with the guanine-binding loops suggests that the effect of the mutation initiating at the Mg^2+^ ion coordination and on to the β-phosphate, propagates through the GDP nucleotide.

The remaining perturbed residues are distant from the GDP nucleotide and can be described as forming two successive layers. The first layer comprises five residues located at the rear of the guanine base-binding residues (Figure 5D). Three of them (S94, Q128, and F124) were destabilized and two (Q156 and A157) were stabilized.

The furthermost interaction layer is comprised of seven residues located in or near helix 4 (residues E102, M108, E113, and R109, Figure 5E) and helix 5 (residues N135, E138, W153, Figure 5F). Three residues in the N-terminus, middle and C-terminus of helix 4 are destabilized, while only one residue (R109) is stabilized. In helix 5, N135 (131 in Arf6 numbering) and E138, both at the beginning of helix 5, are destabilized by the I42S mutation. W153, which engages in CH π edge interactions with H146 just after the kink in helix 5, is also destabilized in the I42S mutant. Taken together, the residues perturbed by the I42S mutation trace a belt from the switch region to the backside of the protein, likely via the Mg^2+^ ion and through the GDP ligand (Figure 5A).

## Discussion

### An allosteric belt extends from the switch region of Arf1 through the GDP ligand to the C-terminal backside

Our HP NMR-based approach has revealed that the I42S mutation, which is in the front of Arf1, perturbs residues along a belt that extends to helices 4 and 5 at the C-terminal backside of Arf1. Remarkably, these residues define a connected pathway that includes the Mg^2+^ ion and the GDP nucleotide. We propose that this pathway delineates an allosteric communication pathway between the front and backside of the Arf proteins. Interestingly, the nucleotide itself plays an integral energetic role in this pathway, which resembles a belt in which the GDP ligand serves as the buckle. It should be noted that the proposed pathway comprises only one residue from the switch region, itself. We cannot rule out effects of the mutation on the stability of residues in the switch since several switch residues near the I42S mutation, 39, 40, 43, 44, and 46 in switch 1 and 51, 52, and 58 in the interswitch, could not be unambiguously assigned in the amide ^1^H-^15^N HSQC spectrum of Arf1 I42S due to their chemical shift perturbations (Table S1). Thus, they could not be studied by our approach.

We previously hypothesized, based on their very high residue-specific stability, that residues in helices 4 and 5 of Arf1 acted as allosteric constraints against switching (27). Residues in these helices were found to be less stable in Arf6 than in Arf1 (26). We then demonstrated via site-directed mutagenesis that the local stability of residues at the beginning of helix 5 on the backside of Arf6 controlled the switching rate on the front side (26). Mutating these residues in helix 5 of Arf6 to their corresponding Arf1 residues led to local stabilization as well as slower switching. In particular, the substitution of KPH 131-133 (135-137 in Arf1 numbering) in Arf6 by the NAA in Arf1 led to the most effective local stabilization and the slowest switching rate. We termed this relationship “back-to-front” allosteric control. Moreover, our prior evolutionary covariance analysis of long-distance covarying residues (26) revealed very high probability covariance involving this region on the back side of the small GTPases across a very large multiple sequence alignment of homologs, suggesting that the energetic networks on the backside of this large family are finely tuned to control switching via the allosteric belt.

This allosteric belt is not the only pathway of communication within the Arf proteins. The N-terminal helix, unique to the Arf subfamily of GTPases, represses switching via interactions with the interswitch (16, 29). Dissociation of the N-terminal helix from the Arf core and its interaction with membranes (including the myristoyl lipid) both favors activation and localizes Arf function to membranes. This effect was described as an allosteric pathway between the N-terminal helix and the switch 1 and 2 regions via the interswitch (29). We showed recently that the entire protein, including the switch region, is significantly destabilized in the Arf1Δ17 mutant (28). This global destabilization upon deletion of the N-terminal helix contrasts starkly with the front-to-back belt of perturbations associated with the I42S mutation. We therefore propose that Arf proteins possess two distinct allosteric communication pathways, which operate via entirely distinct mechanisms. How these pathways cooperate during the GDP/GTP cycle, as Arf proteins interact with diverse membranes and proteins, will be an important question for future studies.

### The allosteric path linking the backside of Arf GTPases to the switch region is bidirectional

Along with our previous demonstration of back-to-front control of switching, the present results demonstrate that the I42S mutation on the front side of the protein likewise leads to perturbations in residue-specific stability on the back side. Thus, in addition to “back-to-front” control of switching, here we observe “front-to-back” energetic communication. Since allosteric communication must be bidirectional, as required by thermodynamics, the present observation of bidirectionality in this allosteric communication pathway is not surprising. Indeed, reciprocal ligand interactions have been amply reported. The oldest and most well-known example may be the Bohr effect in hemoglobin (33), in which proton and oxygen binding are energetically and reciprocally coupled. However, to our knowledge, experimental evidence for two-way energetic coupling within an allosteric network of interacting residues in a protein sequence has not been reported previously.

### The allosteric network in the Arf GTPases controls access to the MG ensemble, rather than controlling the switch itself

The pressure dependence of spontaneous switching in the Δ17 variants of WT Arf1 and the I42S mutant provides insight into the mechanism by which switching is enhanced by pressure and the I42S mutation. We have shown that at intermediate pressures, ∼1500 bar, the Arfs populate a functional MG ensemble that is only a few kcal less stable than the native state. The GDP nucleotide remains bound in this MG ensemble, and the protein remains globular and retains secondary structure. However, the Arfs are dynamically disordered in this MG ensemble, and less stable than in the native state. Populating this excited-state ensemble, even transiently, increases the probability of accessing the even more disrupted states involved in the switch itself. The fact that at high pressure spontaneous switching occurs at identical rates for the WT and mutant indicates that neither pressure nor the I42S mutation enhances the switch transition per se. Instead, both pressure and the I42S mutation favor switching by increasing the probability of populating the MG ensemble from which the actual switch can occur. Hence, once in the MG state, the actual conformational change associated with the switch occurs at the same rate for WT and mutant Arf1.

## Conclusions

In prior work, we established that a functional MG ensemble is a major feature of the conformational landscape of Arf GTPases (27). This broad excited state ensemble plays a central functional role, priming the Arfs for their nucleotide switch. We also revealed and characterized an allosteric network in this protein family that links the C-terminal back of the protein to the switch region (26). Here we map the actual pathway of this allosteric communication pathway and demonstrate that it is bidirectional, as required by thermodynamics. It runs between the switch region and the back of the protein, passing through the GDP ligand. Moreover, we demonstrate that rather than impacting the nucleotide switch transition itself, the allosteric mechanism of this pathway is to modulate the population of the MG ensemble, thereby impacting the overall switching probability. In future work, it will be of interest to determine the extent to which this allosteric network is operative in the broader family of small GTPases (Arf-like/Sar, Ran, Rab, Rho and Ras GTPases) and to establish the energetic and structural implications of their interaction networks for the switching mechanisms of these important proteins.

## Acknowledgments

The work was funded by a grant from the National Institutes of Health, GM 137766 to CAR, and grants from the Fondation pour la Recherche Médicale (grant EQU202003010344) and the French Academy of Sciences (Grand Prix Emile Jungfleich, 2019) to J.C. The Center for High Energy X-ray Science (CHEXS) is supported by the NSF award DMR-2342336, and the MacCHESS resource is supported by NIGMS award 1-P30-GM124166-01A1. The Rensselaer Polytechnic Institute NMR core facility acknowledges National Institute of Standards and Technology grant 60NANB22D167 and National Institute of Health Grant 1S10OD 030482-01.

## Supplemental Information for

**Table S1.**
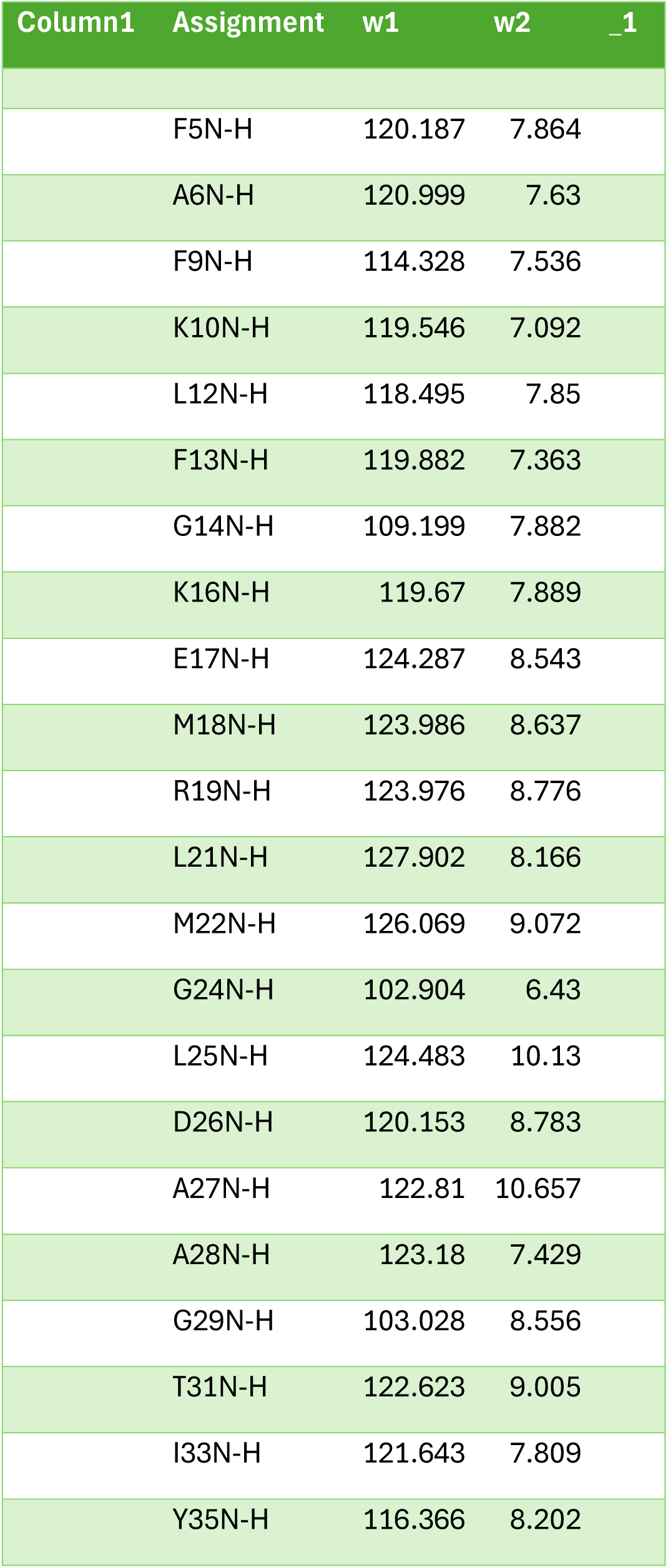
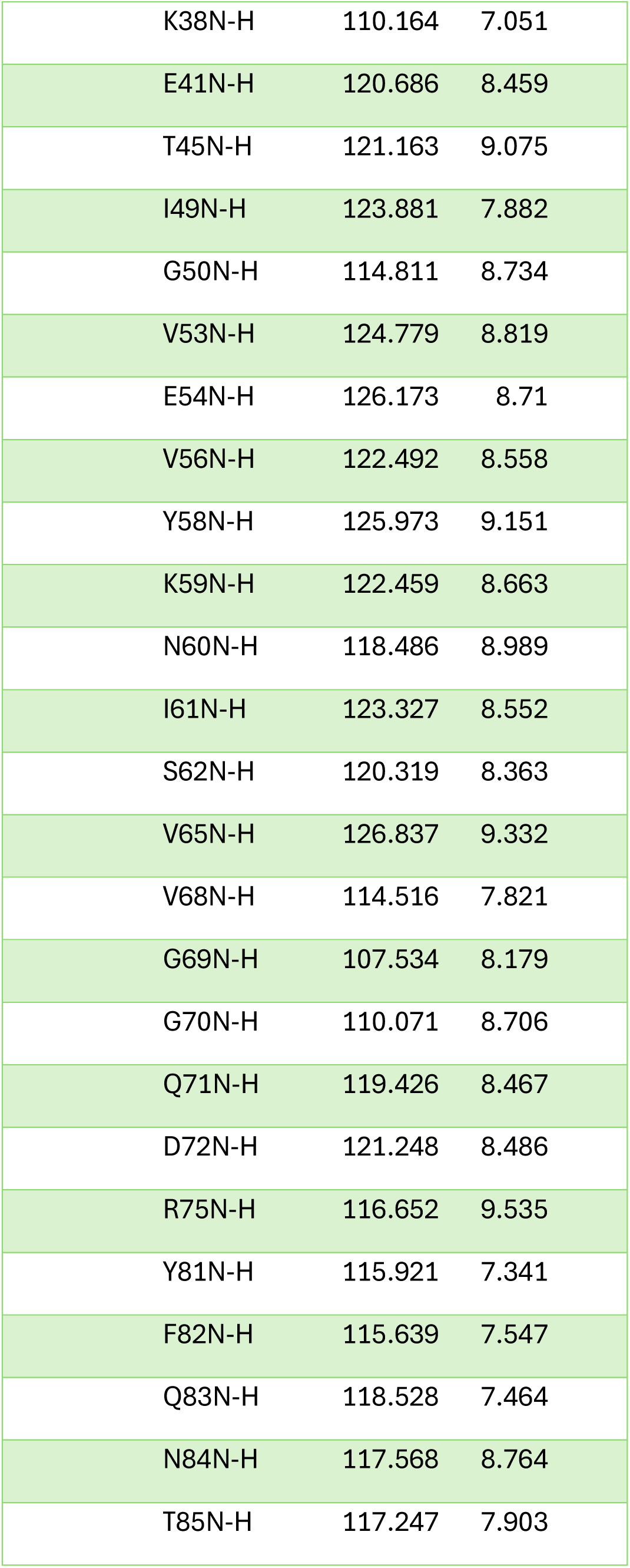
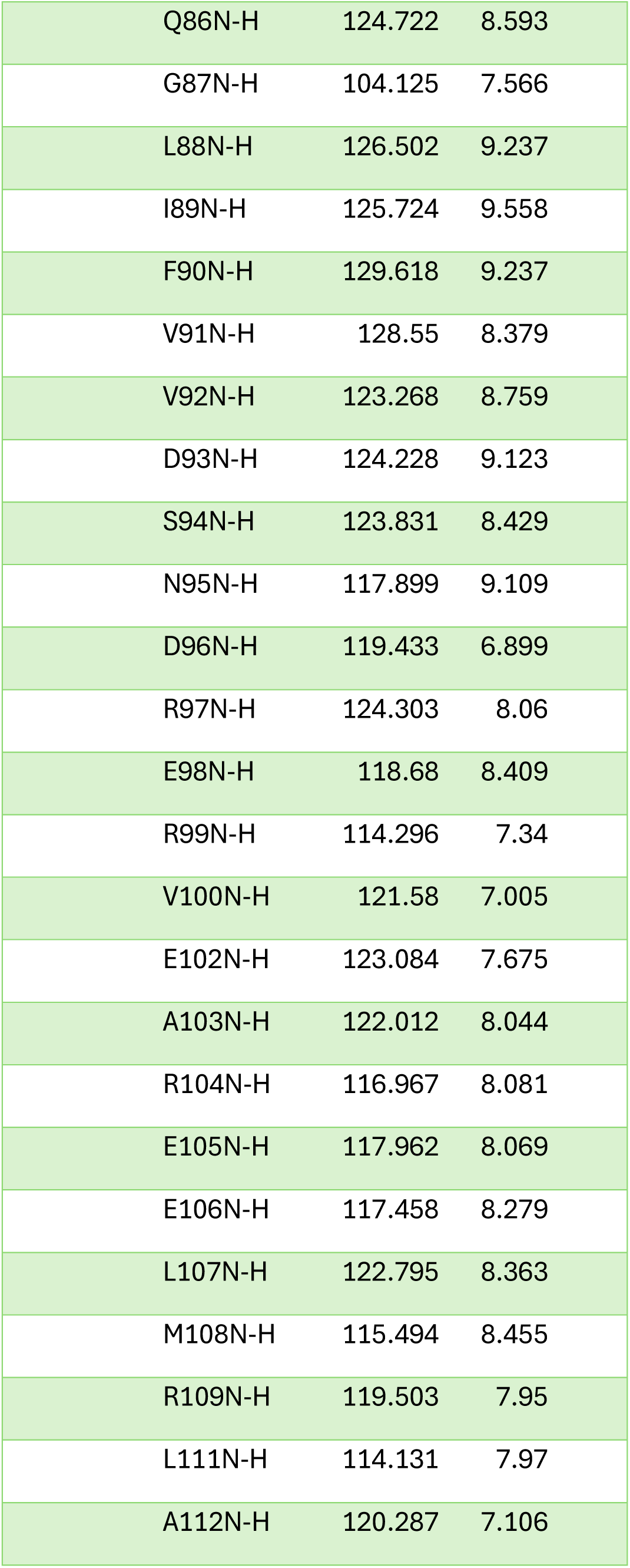
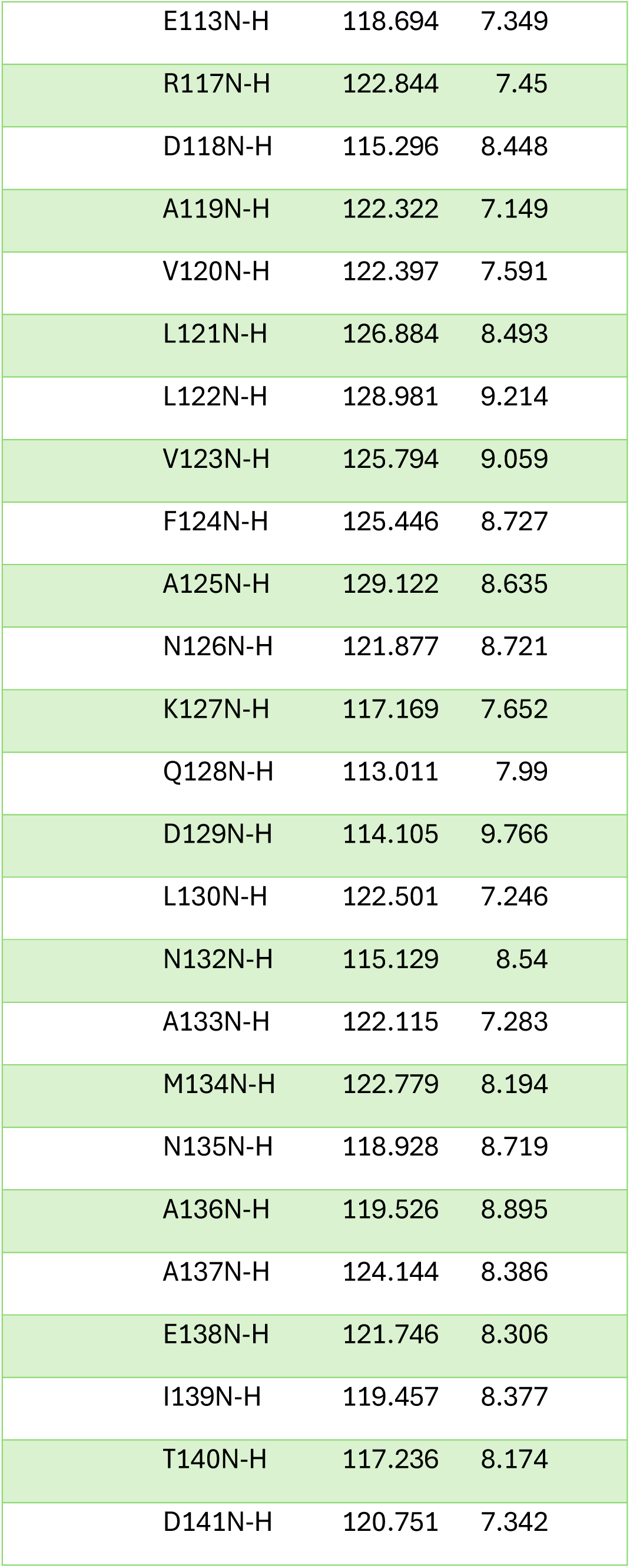
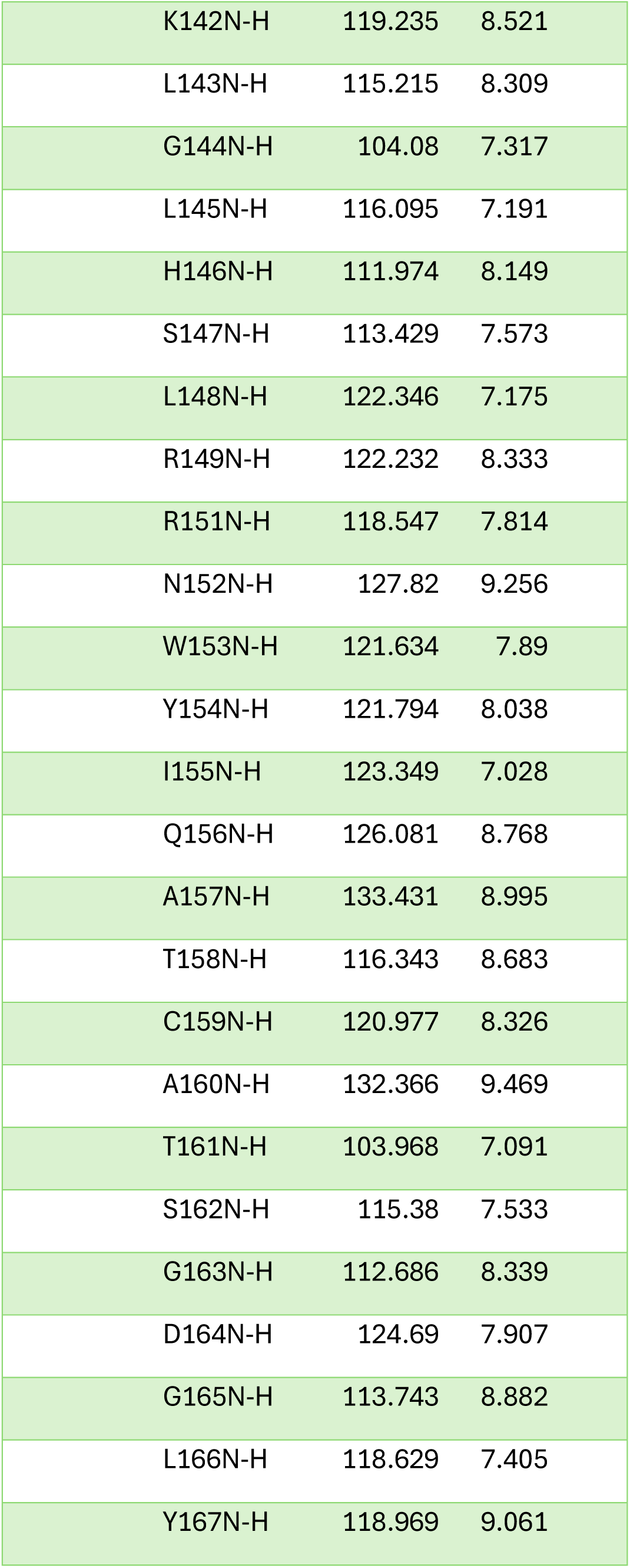
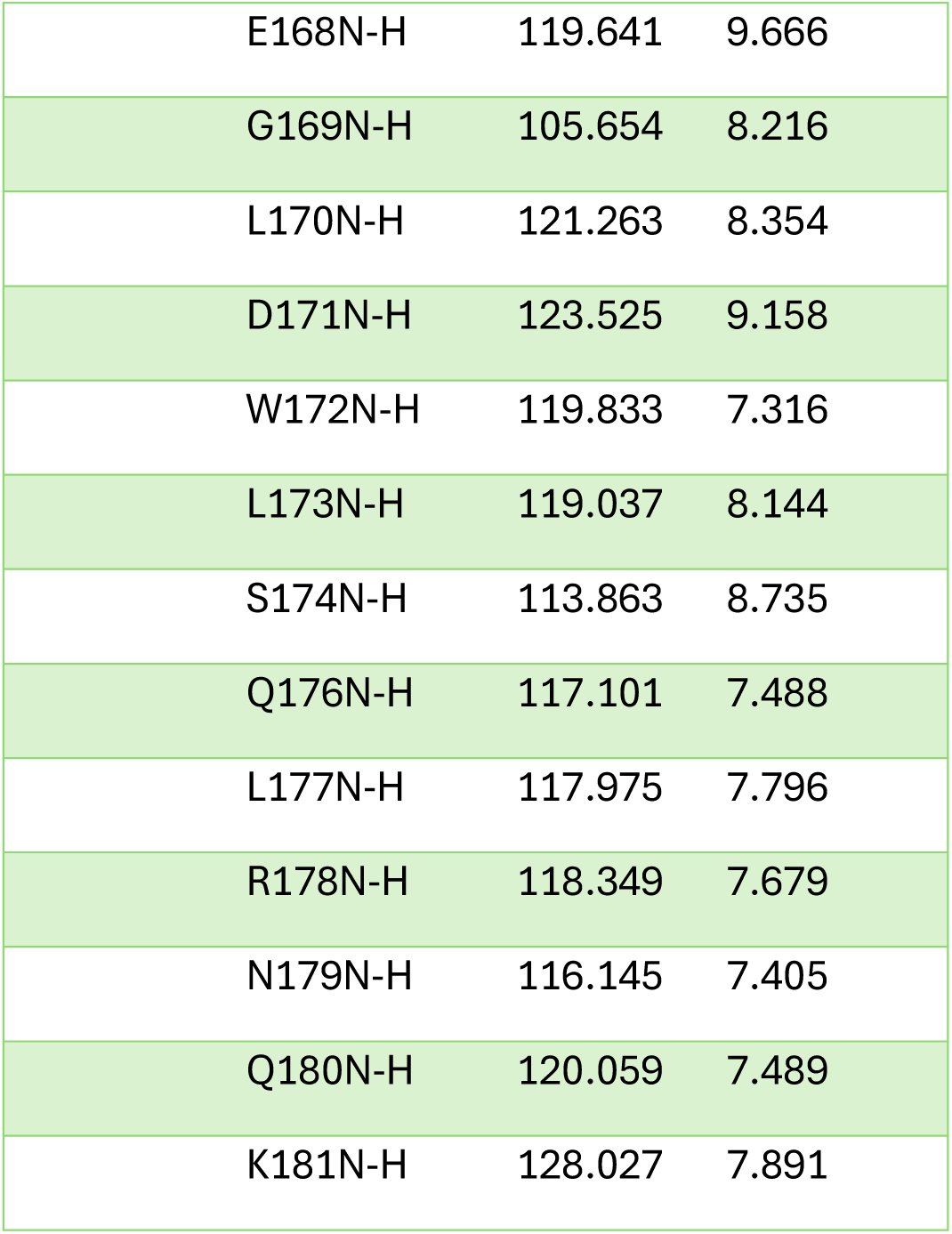
^1^H-^15^N amide backbone assignments for Arf1 I42S.

| Column1 | Assignment | w1 | w2 | _1 |
| --- | --- | --- | --- | --- |
|  | F5N-H | 120.187 | 7.864 |  |
|  | A6N-H | 120.999 | 7.63 |  |
|  | F9N-H | 114.328 | 7.536 |  |
|  | K10N-H | 119.546 | 7.092 |  |
|  | L12N-H | 118.495 | 7.85 |  |
|  | F13N-H | 119.882 | 7.363 |  |
|  | G14N-H | 109.199 | 7.882 |  |
|  | K16N-H | 119.67 | 7.889 |  |
|  | E17N-H | 124.287 | 8.543 |  |
|  | M18N-H | 123.986 | 8.637 |  |
|  | R19N-H | 123.976 | 8.776 |  |
|  | L21N-H | 127.902 | 8.166 |  |
|  | M22N-H | 126.069 | 9.072 |  |
|  | G24N-H | 102.904 | 6.43 |  |
|  | L25N-H | 124.483 | 10.13 |  |
|  | D26N-H | 120.153 | 8.783 |  |
|  | A27N-H | 122.81 | 10.657 |  |
|  | A28N-H | 123.18 | 7.429 |  |
|  | G29N-H | 103.028 | 8.556 |  |
|  | T31N-H | 122.623 | 9.005 |  |
|  | I33N-H | 121.643 | 7.809 |  |
|  | Y35N-H | 116.366 | 8.202 |  |

|  |  |  |
| --- | --- | --- |
| K38N-H | 110.164 | 7.051 |
| E41N-H | 120.686 | 8.459 |
| T45N-H | 121.163 | 9.075 |
| I49N-H | 123.881 | 7.882 |
| G50N-H | 114.811 | 8.734 |
| V53N-H | 124.779 | 8.819 |
| E54N-H | 126.173 | 8.71 |
| V56N-H | 122.492 | 8.558 |
| Y58N-H | 125.973 | 9.151 |
| K59N-H | 122.459 | 8.663 |
| N60N-H | 118.486 | 8.989 |
| I61N-H | 123.327 | 8.552 |
| S62N-H | 120.319 | 8.363 |
| V65N-H | 126.837 | 9.332 |
| V68N-H | 114.516 | 7.821 |
| G69N-H | 107.534 | 8.179 |
| G70N-H | 110.071 | 8.706 |
| Q71N-H | 119.426 | 8.467 |
| D72N-H | 121.248 | 8.486 |
| R75N-H | 116.652 | 9.535 |
| Y81N-H | 115.921 | 7.341 |
| F82N-H | 115.639 | 7.547 |
| Q83N-H | 118.528 | 7.464 |
| N84N-H | 117.568 | 8.764 |
| T85N-H | 117.247 | 7.903 |
| Q86N-H | 124.722 | 8.593 |
| G87N-H | 104.125 | 7.566 |
| L88N-H | 126.502 | 9.237 |
| I89N-H | 125.724 | 9.558 |
| F90N-H | 129.618 | 9.237 |
| V91N-H | 128.55 | 8.379 |
| V92N-H | 123.268 | 8.759 |
| D93N-H | 124.228 | 9.123 |
| S94N-H | 123.831 | 8.429 |
| N95N-H | 117.899 | 9.109 |
| D96N-H | 119.433 | 6.899 |
| R97N-H | 124.303 | 8.06 |
| E98N-H | 118.68 | 8.409 |
| R99N-H | 114.296 | 7.34 |
| V100N-H | 121.58 | 7.005 |
| E102N-H | 123.084 | 7.675 |
| A103N-H | 122.012 | 8.044 |
| R104N-H | 116.967 | 8.081 |
| E105N-H | 117.962 | 8.069 |
| E106N-H | 117.458 | 8.279 |
| L107N-H | 122.795 | 8.363 |
| M108N-H | 115.494 | 8.455 |
| R109N-H | 119.503 | 7.95 |
| L111N-H | 114.131 | 7.97 |
| A112N-H | 120.287 | 7.106 |
| E113N-H | 118.694 | 7.349 |
| R117N-H | 122.844 | 7.45 |
| D118N-H | 115.296 | 8.448 |
| A119N-H | 122.322 | 7.149 |
| V120N-H | 122.397 | 7.591 |
| L121N-H | 126.884 | 8.493 |
| L122N-H | 128.981 | 9.214 |
| V123N-H | 125.794 | 9.059 |
| F124N-H | 125.446 | 8.727 |
| A125N-H | 129.122 | 8.635 |
| N126N-H | 121.877 | 8.721 |
| K127N-H | 117.169 | 7.652 |
| Q128N-H | 113.011 | 7.99 |
| D129N-H | 114.105 | 9.766 |
| L130N-H | 122.501 | 7.246 |
| N132N-H | 115.129 | 8.54 |
| A133N-H | 122.115 | 7.283 |
| M134N-H | 122.779 | 8.194 |
| N135N-H | 118.928 | 8.719 |
| A136N-H | 119.526 | 8.895 |
| A137N-H | 124.144 | 8.386 |
| E138N-H | 121.746 | 8.306 |
| I139N-H | 119.457 | 8.377 |
| T140N-H | 117.236 | 8.174 |
| D141N-H | 120.751 | 7.342 |
| K142N-H | 119.235 | 8.521 |
| L143N-H | 115.215 | 8.309 |
| G144N-H | 104.08 | 7.317 |
| L145N-H | 116.095 | 7.191 |
| H146N-H | 111.974 | 8.149 |
| S147N-H | 113.429 | 7.573 |
| L148N-H | 122.346 | 7.175 |
| R149N-H | 122.232 | 8.333 |
| R151N-H | 118.547 | 7.814 |
| N152N-H | 127.82 | 9.256 |
| W153N-H | 121.634 | 7.89 |
| Y154N-H | 121.794 | 8.038 |
| I155N-H | 123.349 | 7.028 |
| Q156N-H | 126.081 | 8.768 |
| A157N-H | 133.431 | 8.995 |
| T158N-H | 116.343 | 8.683 |
| C159N-H | 120.977 | 8.326 |
| A160N-H | 132.366 | 9.469 |
| T161N-H | 103.968 | 7.091 |
| S162N-H | 115.38 | 7.533 |
| G163N-H | 112.686 | 8.339 |
| D164N-H | 124.69 | 7.907 |
| G165N-H | 113.743 | 8.882 |
| L166N-H | 118.629 | 7.405 |
| Y167N-H | 118.969 | 9.061 |
| E168N-H | 119.641 | 9.666 |
| G169N-H | 105.654 | 8.216 |
| L170N-H | 121.263 | 8.354 |
| D171N-H | 123.525 | 9.158 |
| W172N-H | 119.833 | 7.316 |
| L173N-H | 119.037 | 8.144 |
| S174N-H | 113.863 | 8.735 |
| Q176N-H | 117.101 | 7.488 |
| L177N-H | 117.975 | 7.796 |
| R178N-H | 118.349 | 7.679 |
| N179N-H | 116.145 | 7.405 |
| Q180N-H | 120.059 | 7.489 |
| K181N-H | 128.027 | 7.891 |

**Table S2.** Parameters from fits of the HP NMR peak intensities vs. pressure for Arf1I42S.

| I42S FLArf1 pH 6.5 |  |  |  |  |
| --- | --- | --- | --- | --- |
| Residue # | $\Delta G_{app}$ | $\sigma_{\Delta G}$ | $\Delta V_{app}$ | $\sigma_{\Delta V}$ |
| 5.00 | 1.63 | 0.13 | 58.84 | 5.84 |
| 6.00 | 2.05 | 0.20 | 42.56 | 7.67 |
| 9.00 | 2.18 | 0.15 | 63.63 | 5.96 |
| 10.00 | 1.64 | 0.11 | 53.63 | 4.46 |
| 12.00 | 1.35 | 0.21 | 47.96 | 10.54 |
| 13.00 | 1.38 | 0.18 | 52.22 | 8.68 |
| 14.00 | 1.47 | 0.08 | 57.02 | 3.44 |
| 16.00 | 1.68 | 0.20 | 39.41 | 8.26 |
| 17.00 | 3.86 | 0.28 | 110.66 | 13.22 |
| 18.00 | 1.32 | 0.12 | 40.30 | 4.80 |
| 19.00 | 2.13 | 0.28 | 59.50 | 13.20 |
| 21.00 | 1.66 | 0.16 | 64.59 | 7.86 |
| 24.00 | 1.63 | 0.13 | 103.19 | 9.31 |
| 25.00 | 2.00 | 0.06 | 57.00 | 2.06 |
| 26.00 | 1.90 | 0.09 | 72.90 | 3.82 |
| 27.00 | 2.21 | 0.23 | 60.03 | 9.81 |
| 28.00 | 4.74 | 0.31 | 119.40 | 13.57 |
| 29.00 | 1.40 | 0.15 | 49.88 | 6.41 |
| 31.00 | 1.02 | 0.19 | 43.95 | 9.95 |
| 33.00 | 1.63 | 0.12 | 57.37 | 4.93 |
| 35.00 | 2.58 | 0.21 | 71.85 | 8.91 |
| 38.00 | 1.31 | 0.20 | 30.63 | 8.24 |
| 41.00 | 1.03 | 0.25 | 39.22 | 13.80 |
| 45.00 | 1.37 | 0.19 | 97.95 | 17.34 |
| 49.00 | 1.09 | 0.19 | 37.99 | 8.82 |
| 53.00 | 2.56 | 0.42 | 67.06 | 24.73 |
| 54.00 | 2.16 | 0.20 | 72.02 | 9.14 |
| 56.00 | 0.71 | 0.21 | 28.54 | 10.96 |
| 58.00 | 2.60 | 0.31 | 83.68 | 17.23 |
| 59.00 | 2.86 | 0.43 | 64.30 | 24.61 |
| 60.00 | 2.07 | 0.17 | 61.20 | 6.92 |
| 61.00 | 0.83 | 0.23 | 41.70 | 14.59 |
| 65.00 | 4.78 | 0.59 | 134.20 | 53.74 |
| 68.00 | 2.79 | 0.23 | 77.81 | 9.93 |
| 69.00 | 1.34 | 0.16 | 62.79 | 9.06 |
| 70.00 | 2.36 | 0.23 | 70.67 | 10.24 |
| 71.00 | 2.07 | 0.22 | 57.59 | 9.27 |
| 72.00 | 1.21 | 0.14 | 51.87 | 7.10 |
| 75.00 | 2.14 | 0.17 | 61.93 | 6.98 |
| 81.00 | 1.53 | 0.09 | 72.04 | 4.52 |
| 84.00 | 1.14 | 0.19 | 83.69 | 16.53 |
| 85.00 | 1.46 | 0.15 | 43.28 | 5.87 |
| 86.00 | 1.96 | 0.17 | 45.87 | 6.31 |
| 87.00 | 1.89 | 0.25 | 45.65 | 10.72 |
| 88.00 | 2.01 | 0.20 | 65.27 | 9.20 |
| 89.00 | 1.92 | 0.13 | 55.37 | 4.82 |
| 90.00 | 1.12 | 0.14 | 44.48 | 6.35 |
| 91.00 | 1.40 | 0.17 | 35.15 | 6.81 |
| 92.00 | 4.18 | 0.44 | 90.22 | 23.47 |
| 93.00 | 1.89 | 0.19 | 56.79 | 7.88 |
| 94.00 | 1.22 | 0.14 | 26.35 | 5.35 |
| 95.00 | 2.11 | 0.07 | 40.50 | 1.94 |
| 96.00 | 1.38 | 0.17 | 24.64 | 6.75 |
| 97.00 | 1.53 | 0.23 | 35.58 | 9.82 |
| 98.00 | 1.25 | 0.12 | 31.80 | 4.53 |
| 99.00 | 1.79 | 0.17 | 33.37 | 6.15 |
| 100.00 | 1.57 | 0.16 | 30.65 | 5.86 |
| 102.00 | 1.61 | 0.16 | 28.37 | 5.78 |
| 103.00 | 2.00 | 0.16 | 48.67 | 5.98 |
| 104.00 | 2.69 | 0.34 | 54.90 | 15.76 |
| 105.00 | 1.61 | 0.15 | 34.62 | 5.49 |
| 106.00 | 2.33 | 0.23 | 64.77 | 10.12 |
| 107.00 | 3.17 | 0.37 | 69.56 | 17.76 |
| 108.00 | 1.46 | 0.20 | 80.11 | 14.23 |
| 109.00 | 3.67 | 0.53 | 76.01 | 34.36 |
| 111.00 | 1.78 | 0.18 | 39.34 | 6.95 |
| 112.00 | 2.21 | 0.11 | 57.78 | 3.62 |
| 113.00 | 1.83 | 0.17 | 43.51 | 6.27 |
| 117.00 | 2.29 | 0.22 | 52.53 | 8.48 |
| 118.00 | 2.07 | 0.23 | 46.64 | 9.03 |
| 119.00 | 3.37 | 0.49 | 64.45 | 29.83 |
| 120.00 | 1.46 | 0.12 | 48.18 | 4.97 |
| 121.00 | 2.99 | 0.43 | 76.04 | 25.47 |
| 122.00 | 1.68 | 0.18 | 26.43 | 6.72 |
| 123.00 | 2.05 | 0.36 | 41.23 | 18.05 |
| 124.00 | 2.09 | 0.20 | 58.78 | 8.09 |
| 125.00 | 1.43 | 0.19 | 27.30 | 7.74 |
| 126.00 | 2.85 | 0.51 | 56.42 | 32.94 |
| 127.00 | 1.82 | 0.19 | 39.88 | 7.21 |
| 128.00 | 1.51 | 0.10 | 32.51 | 3.30 |
| 129.00 | 4.72 | 0.56 | 84.53 | 37.13 |
| 130.00 | 1.92 | 0.28 | 31.94 | 12.49 |
| 132.00 | 1.62 | 0.17 | 32.08 | 6.19 |
| 133.00 | 1.44 | 0.20 | 22.46 | 8.39 |
| 134.00 | 2.78 | 0.35 | 52.35 | 16.62 |
| 135.00 | 1.90 | 0.43 | 23.45 | 26.45 |
| 136.00 | 1.86 | 0.11 | 26.66 | 3.65 |
| 137.00 | 2.15 | 0.32 | 42.90 | 15.01 |
| 138.00 | 1.77 | 0.20 | 28.46 | 7.65 |
| 139.00 | 2.90 | 0.42 | 54.12 | 22.77 |
| 140.00 | 1.73 | 0.21 | 28.54 | 8.13 |
| 141.00 | 1.89 | 0.23 | 35.13 | 9.45 |
| 142.00 | 3.95 | 0.54 | 78.56 | 35.82 |
| 143.00 | 2.37 | 0.44 | 35.17 | 25.76 |
| 144.00 | 2.10 | 0.25 | 33.20 | 10.28 |
| 145.00 | 1.78 | 0.21 | 25.58 | 8.42 |
| 146.00 | 1.54 | 0.25 | 19.55 | 11.42 |
| 148.00 | 2.04 | 0.24 | 37.48 | 9.63 |
| 149.00 | 1.58 | 0.14 | 44.38 | 5.31 |
| 151.00 | 1.49 | 0.16 | 37.35 | 6.27 |
| 152.00 | 3.32 | 0.43 | 63.91 | 23.25 |
| 153.00 | 1.42 | 0.17 | 29.15 | 6.58 |
| 154.00 | 2.00 | 0.21 | 41.10 | 8.16 |
| 155.00 | 2.58 | 0.44 | 46.02 | 25.16 |
| 156.00 | 3.34 | 0.51 | 63.87 | 32.70 |
| 157.00 | 3.21 | 0.51 | 61.10 | 32.78 |
| 158.00 | 2.26 | 0.29 | 43.77 | 12.84 |
| 159.00 | 6.03 | 0.47 | 127.16 | 25.88 |
| 160.00 | 1.65 | 0.16 | 41.47 | 6.13 |
| 161.00 | 1.37 | 0.14 | 29.42 | 5.33 |
| 162.00 | 1.40 | 0.27 | 27.48 | 12.62 |
| 163.00 | 1.89 | 0.21 | 42.89 | 8.30 |
| 164.00 | 1.67 | 0.18 | 33.49 | 6.82 |
| 165.00 | 2.13 | 0.36 | 41.08 | 17.91 |
| 166.00 | 2.11 | 0.16 | 49.28 | 5.89 |
| 167.00 | 1.63 | 0.10 | 53.44 | 3.96 |
| 168.00 | 2.49 | 0.29 | 56.88 | 13.00 |
| 169.00 | 2.17 | 0.32 | 42.37 | 14.66 |
| 170.00 | 1.88 | 0.13 | 49.74 | 4.86 |
| 171.00 | 1.60 | 0.11 | 40.54 | 3.87 |
| 172.00 | 1.39 | 0.15 | 30.85 | 5.67 |
| 173.00 | 2.35 | 0.26 | 56.96 | 11.07 |
| 174.00 | 1.26 | 0.14 | 31.23 | 5.17 |
| 176.00 | 1.80 | 0.19 | 36.99 | 7.30 |
| 177.00 | 2.11 | 0.24 | 47.74 | 10.08 |
| 178.00 | 1.58 | 0.20 | 36.85 | 8.01 |
| 179.00 | 1.87 | 0.28 | 41.55 | 12.77 |
| 180.00 | 1.90 | 0.28 | 43.80 | 12.75 |
| 181.00 | 2.58 | 0.41 | 52.80 | 22.61 |

**Table S3.**
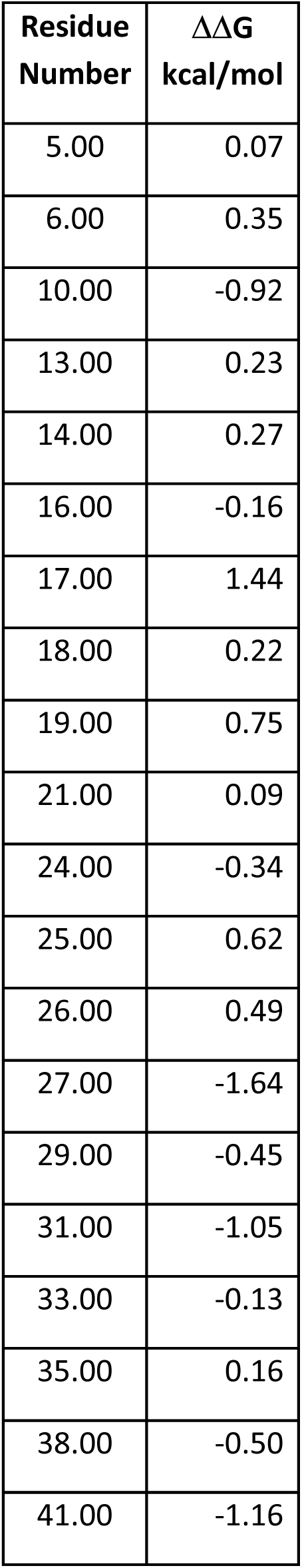
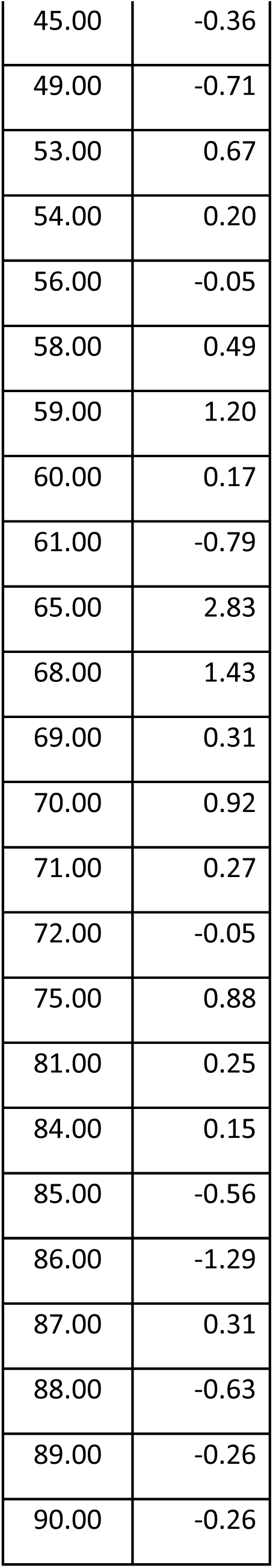
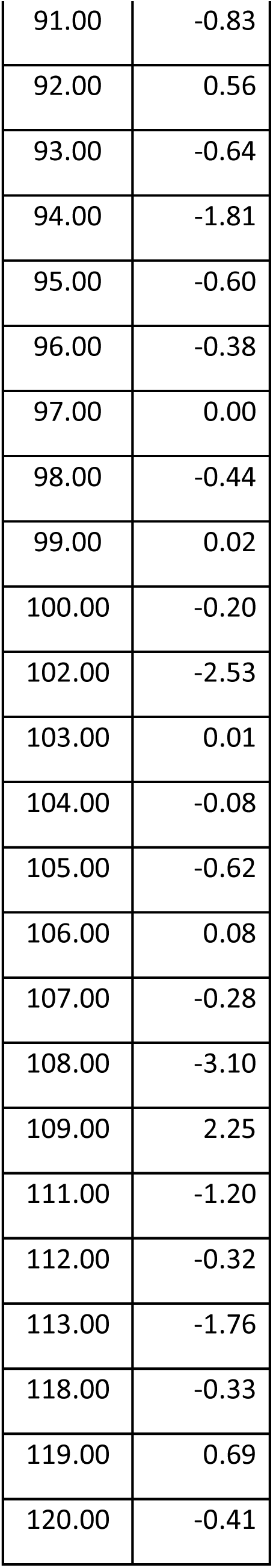
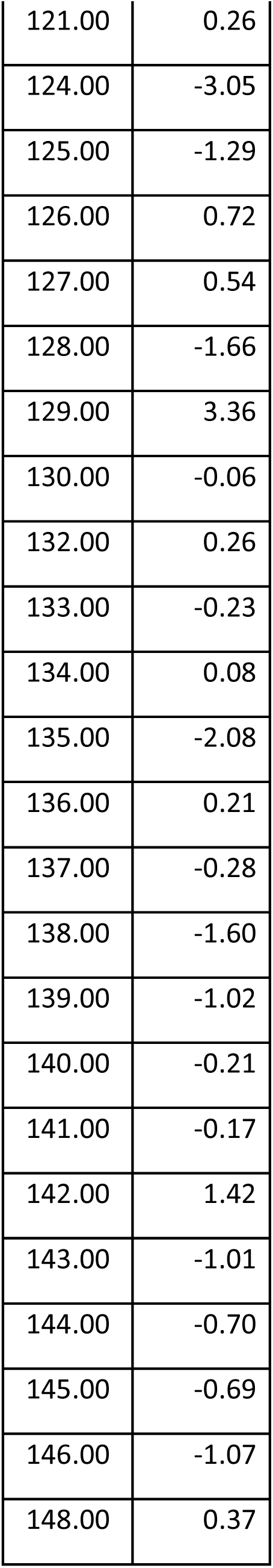
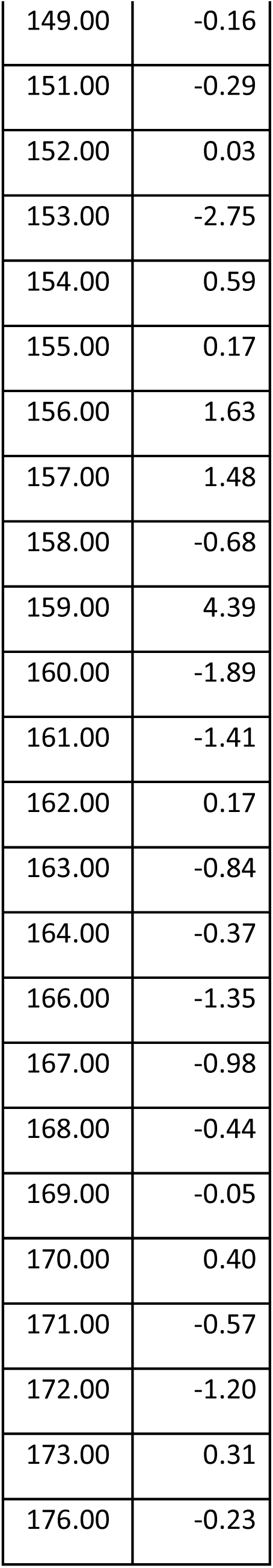
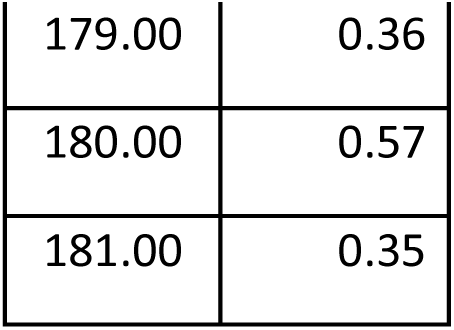
Difference in residue-specific stability between Arf1I42S and WT Arf1.

**Figure S1.**
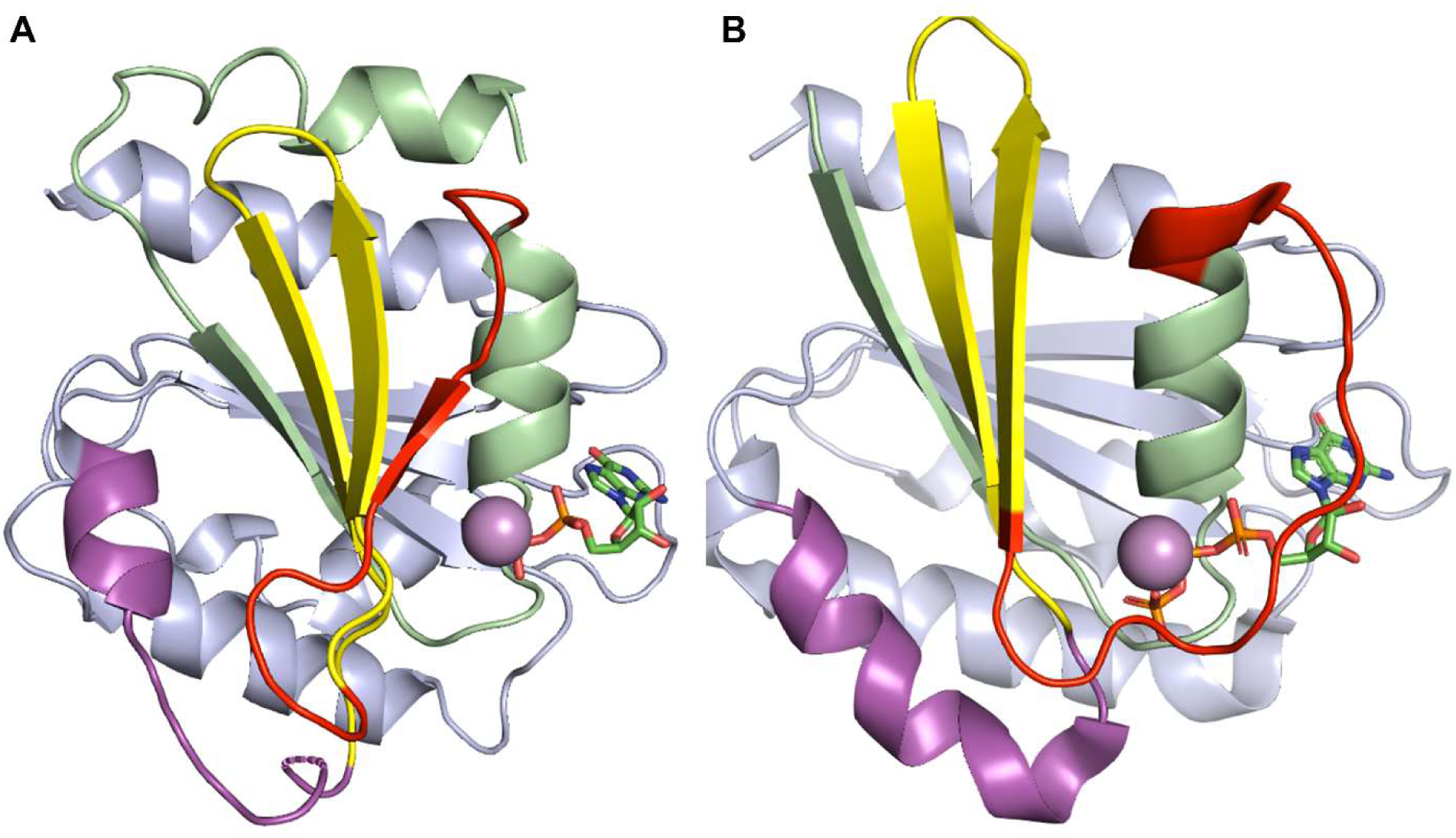
Structural aspects of the nucleotide switch in the Arf proteins. A) The Arf1 GDP-bound structure (pdb code: 1hur from Amor et al. 1994. B) The Arf1 GDP-bound structure (pdb code: 1j2j from Shiba et al 2003. The structure of the GGA1 GAT Nt peptide in 1j2j is not represented. C) Overlay of the Arf1 GDP structure (1hur) with the Arf6-GDP structure, pdb code:1eos from Menetrey et al, 2000. The structures are colored green for the N-terminal region through helix 2, red for Switch 1, yellow for the interswitch and magenta for Switch 2. The rest of the protein is beige.

**Figure S2.**
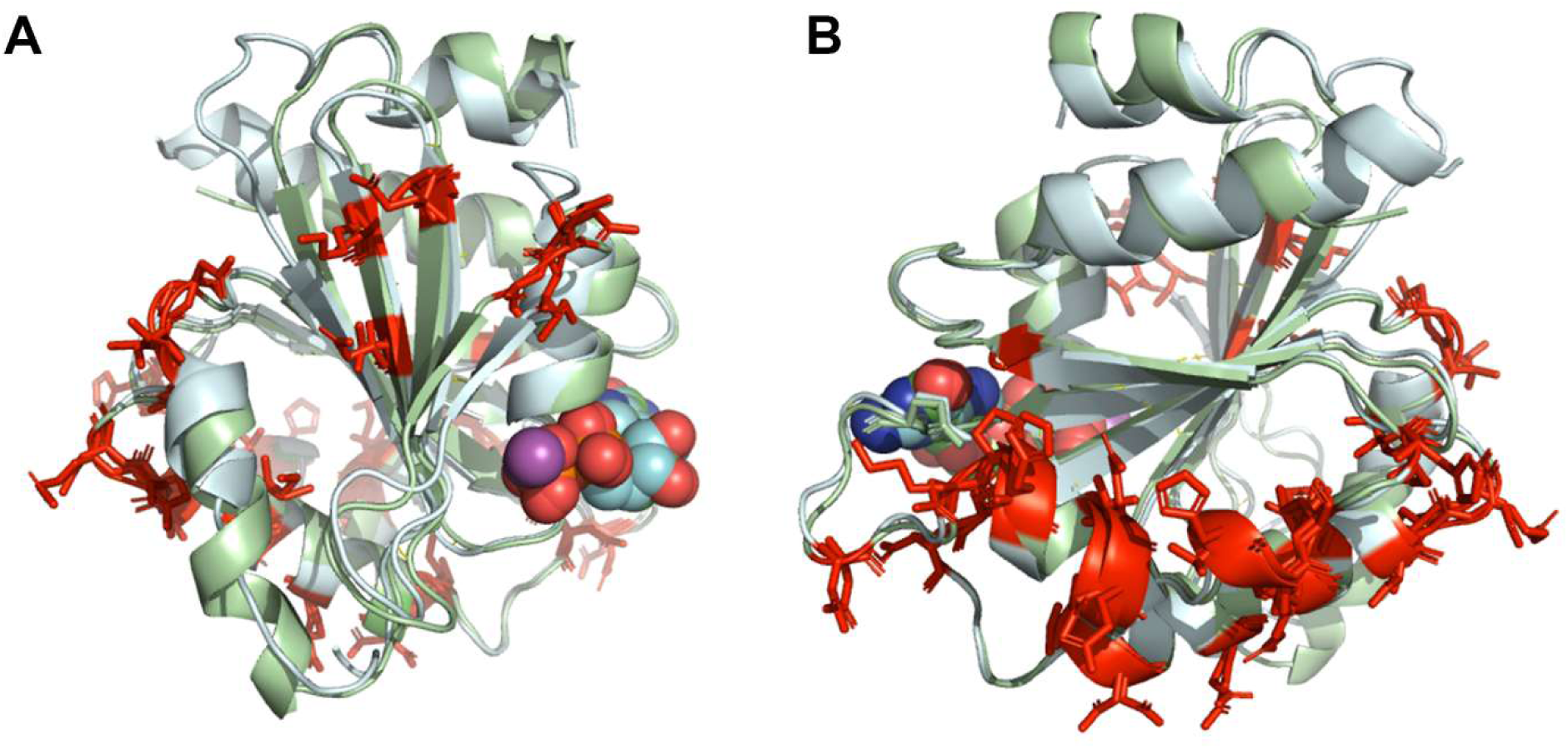
Sequence difference between Arf homologs Arf1 and Arf6. Overlay of Arf1 (light blue) and Arf6 (light green) structures from pdb 1hur and 1eos respectively. Positions corresponding to sequence differences are shown in red sticks. The GDP ligand is shown as CPK spheres. A Front view of the switch region. B) Back view of the C-terminal half of the protein.

**Figure S3.**
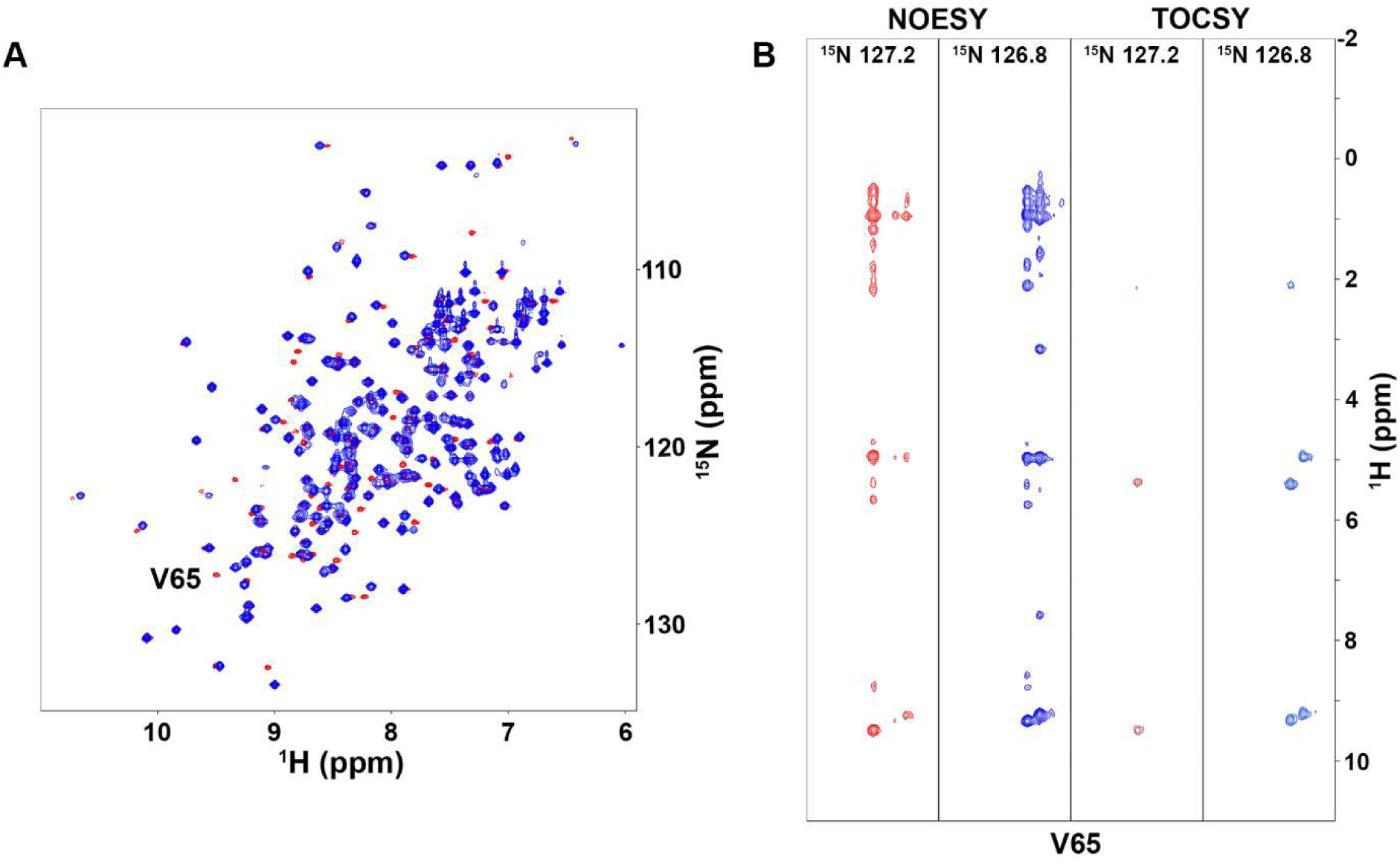
Comparison of the ^1^H-^15^N NMR spectra of WT Arf1 (red) and the I42S variant (blue). A) ^1^H-^15^N HSQC spectra. A few residues exhibit significant chemical shift perturbations due to the I42S mutation. As an example, V65 is highlighted. B) Assignments were verified using ^15^N NOESY and TOCSY experiments. Again, the V65 example is shown.

**Figure S4.**
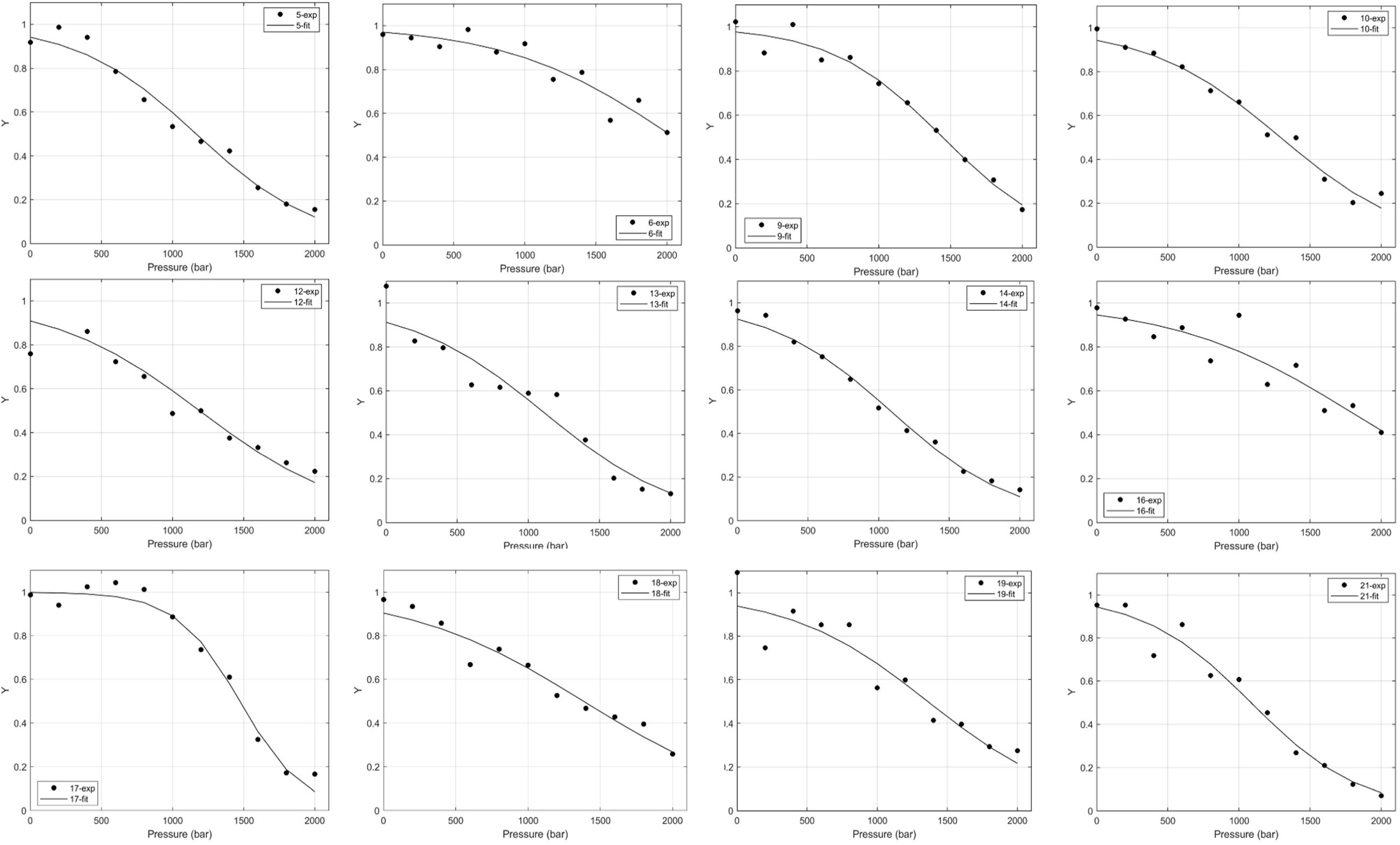
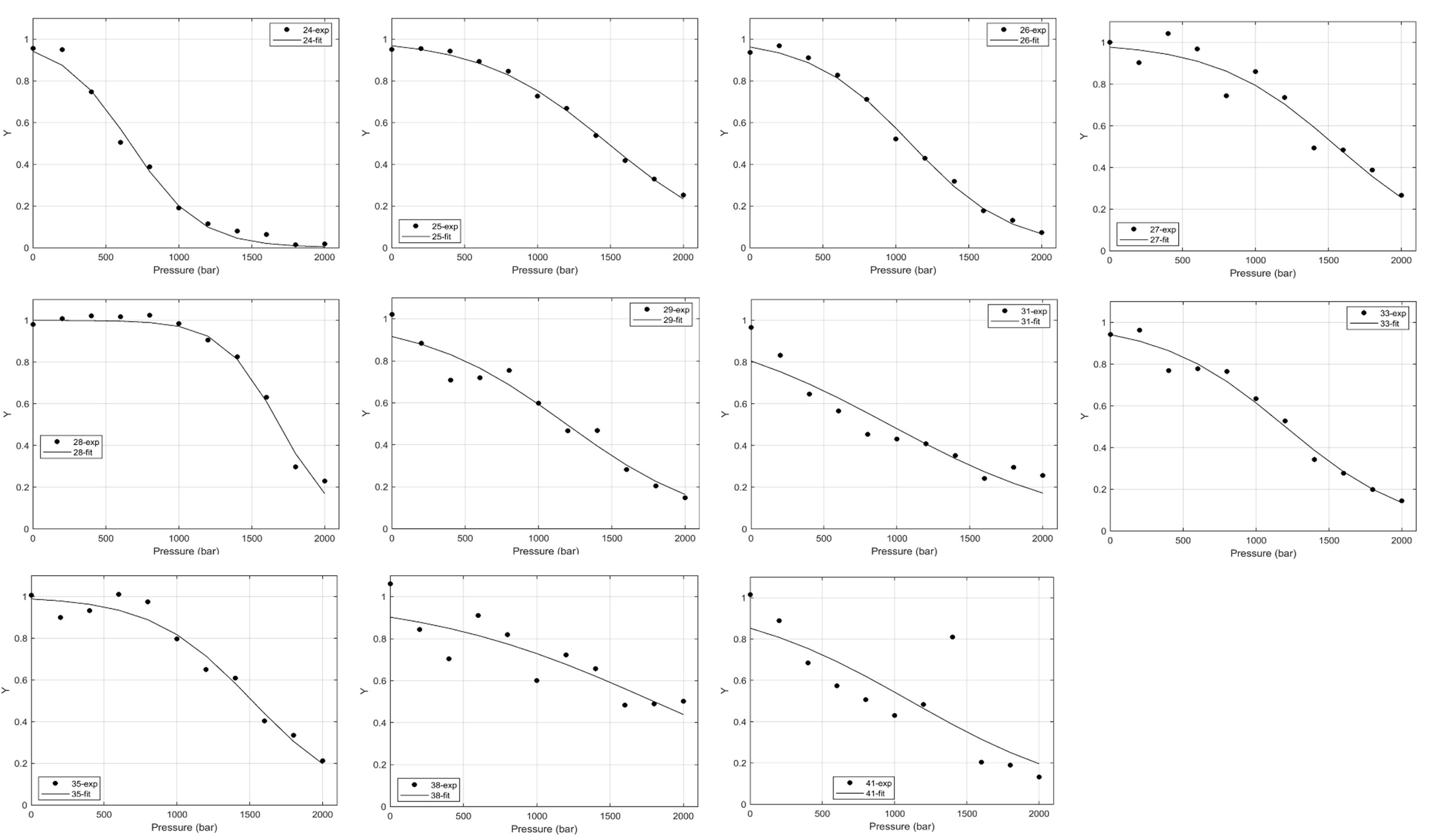
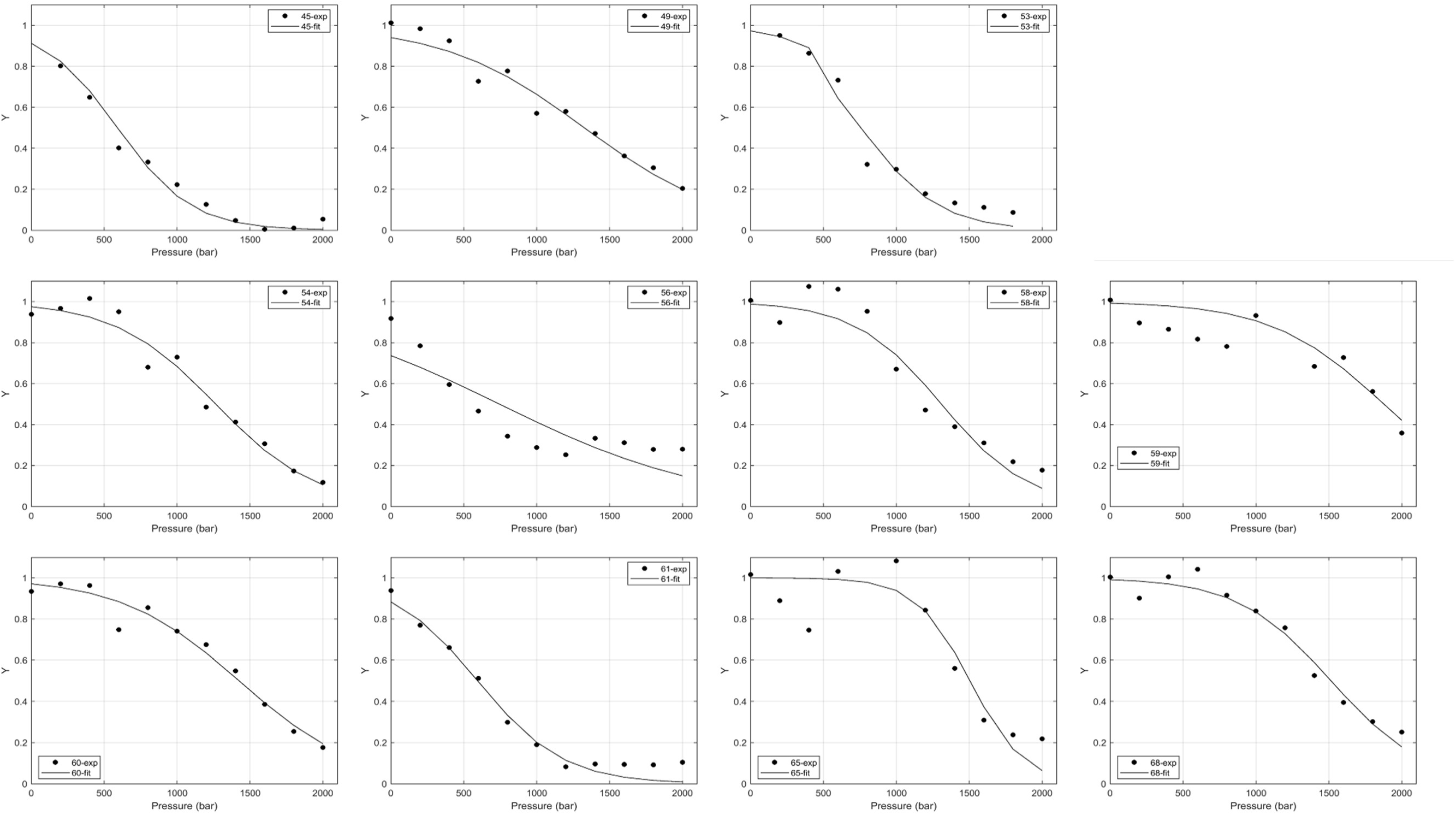
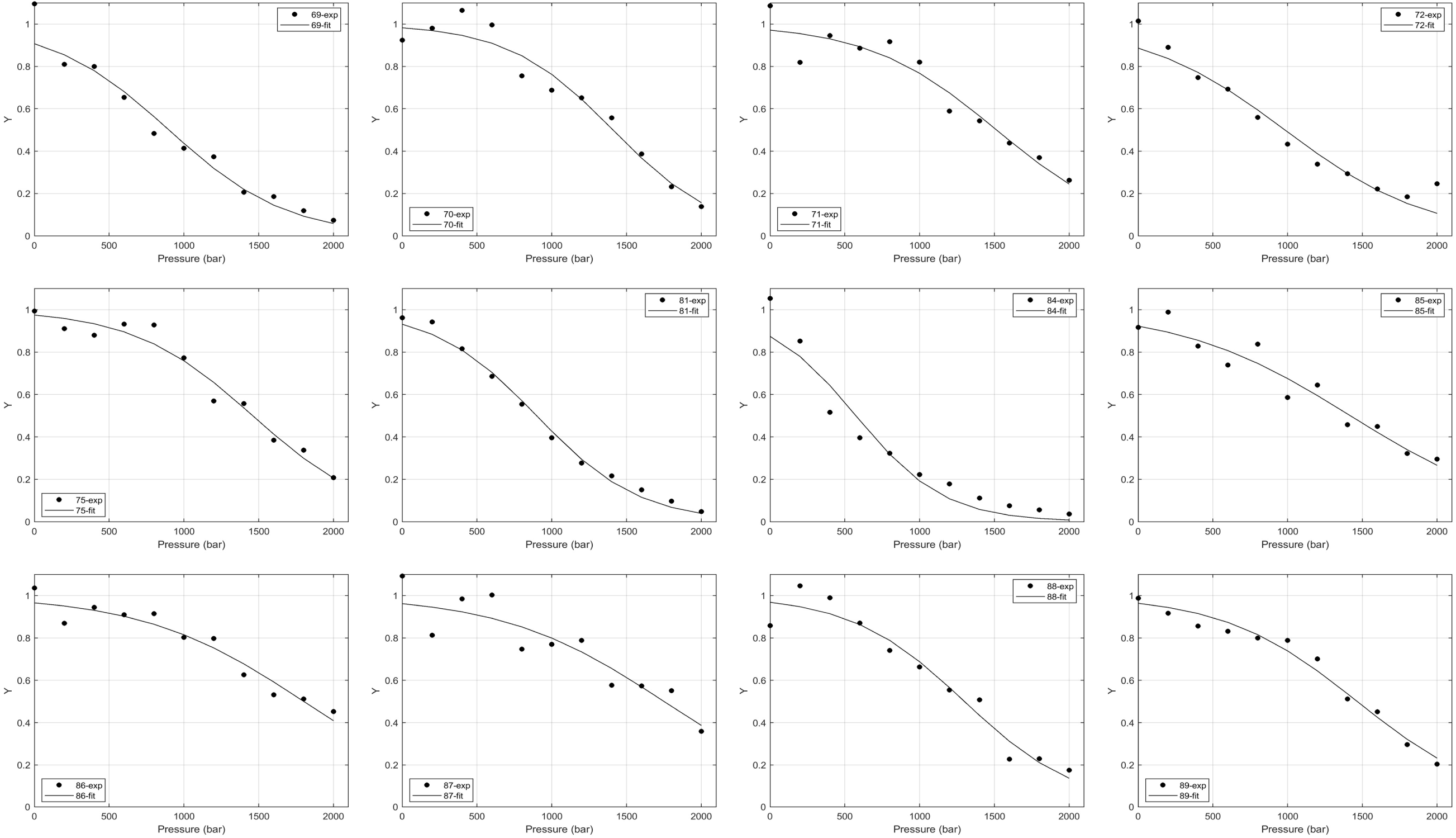
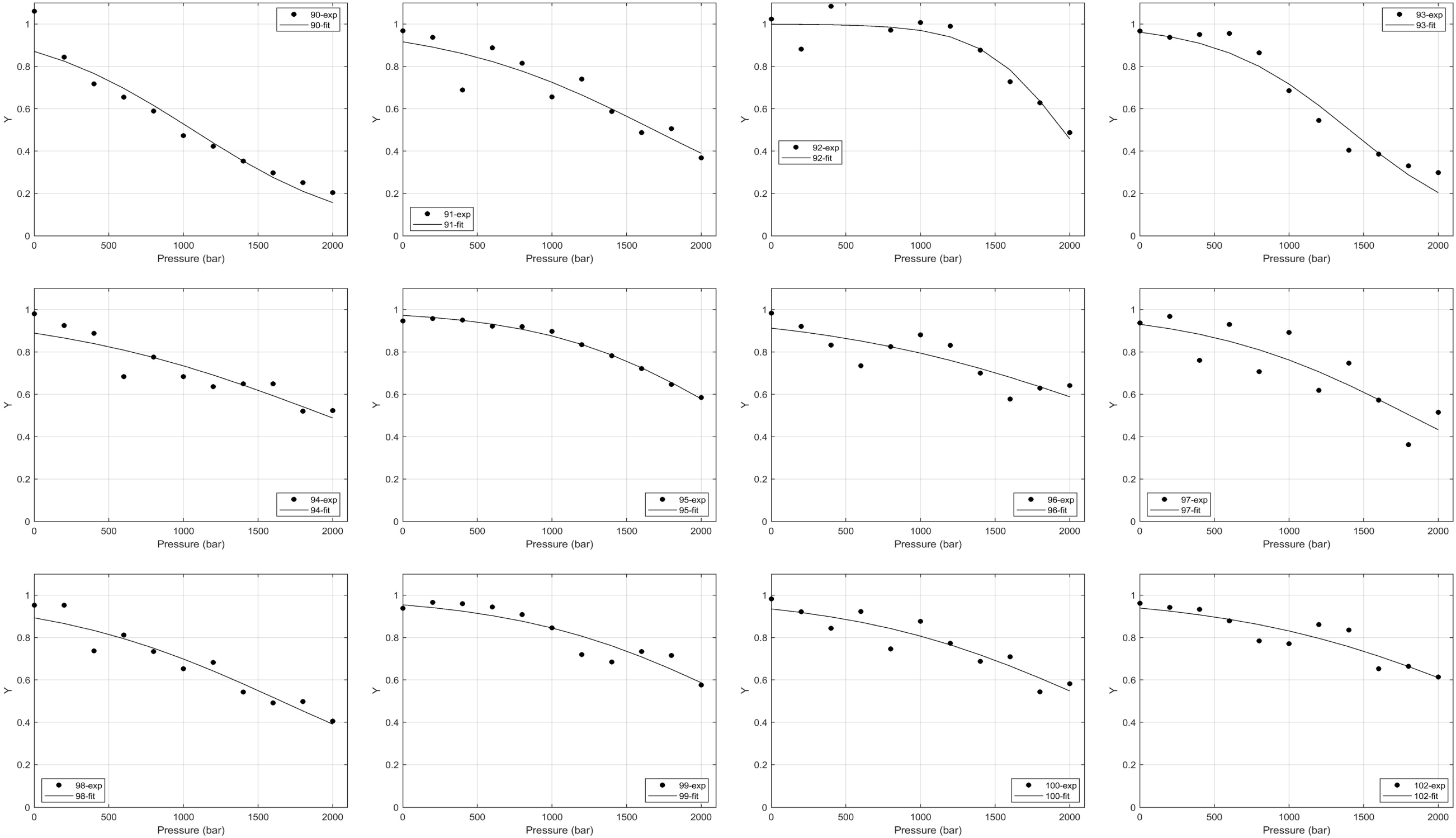
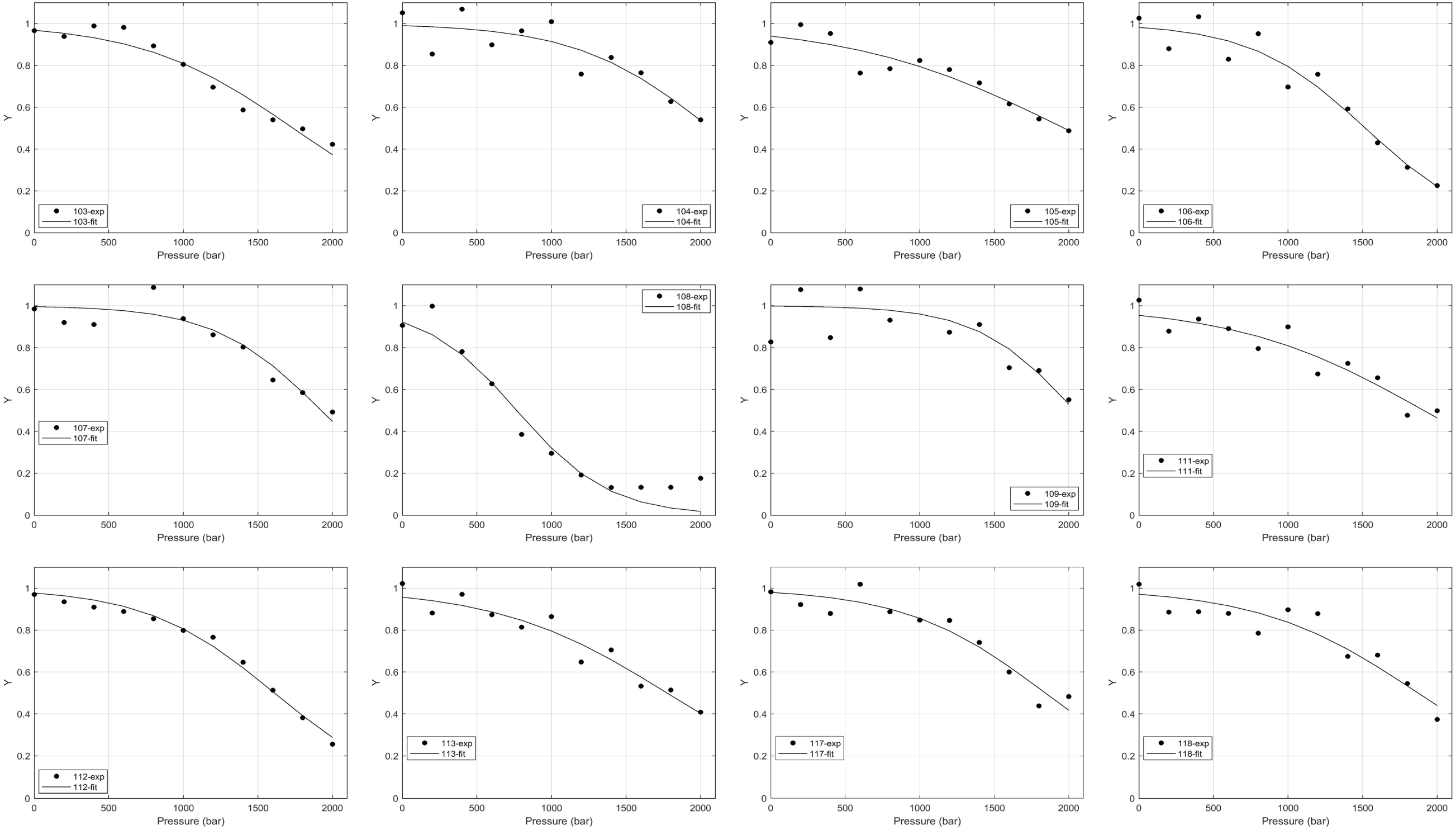
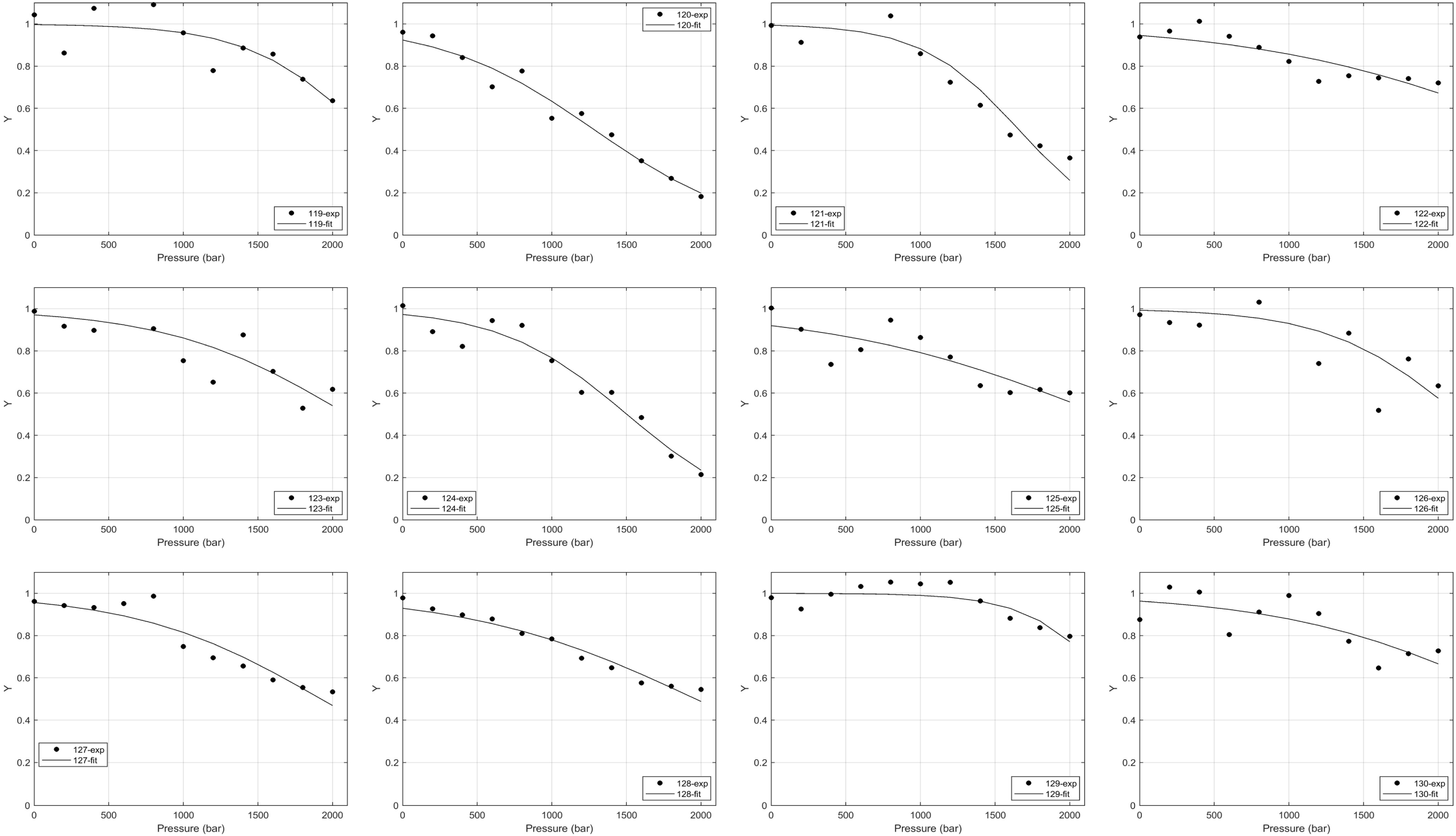
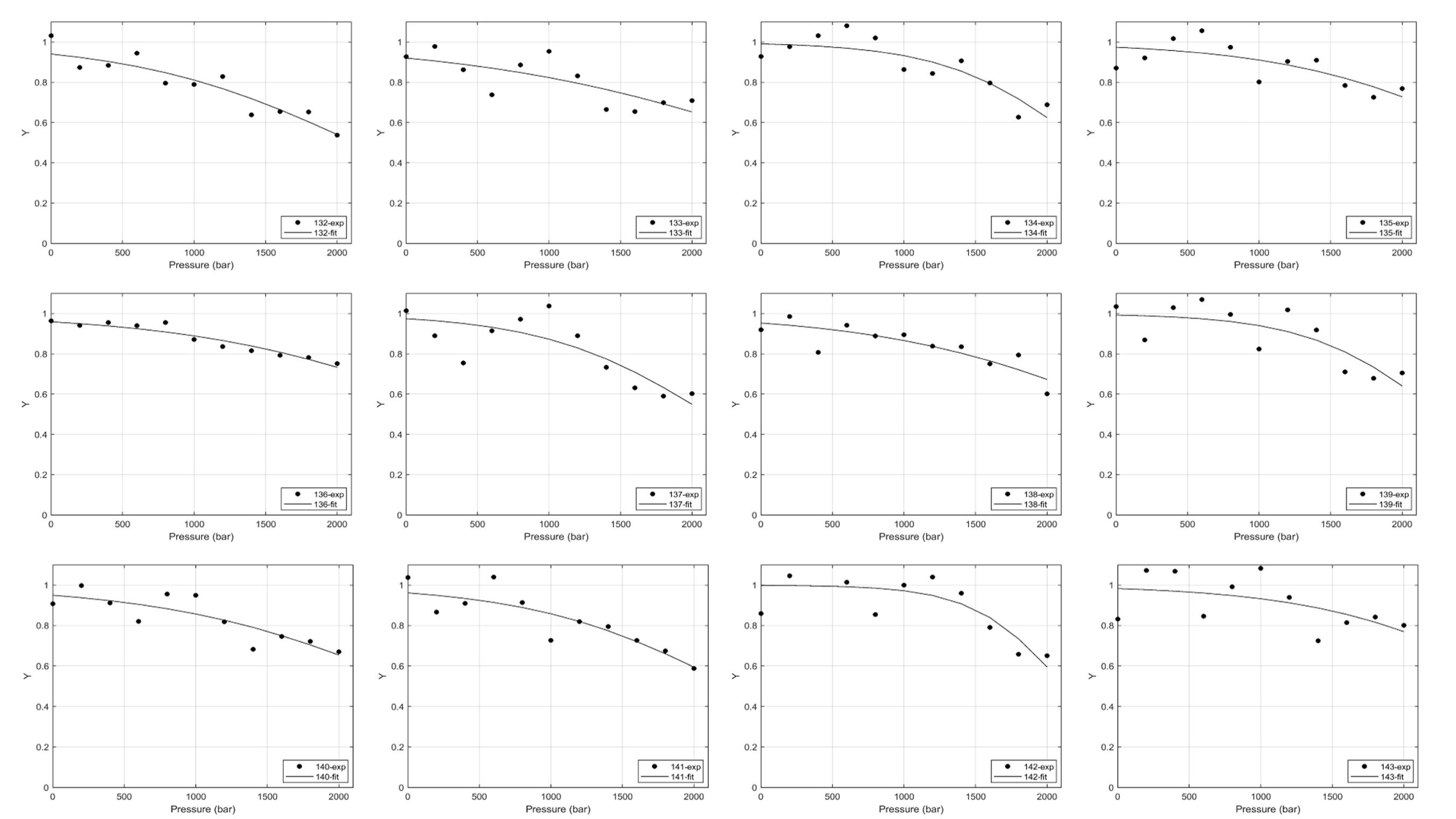
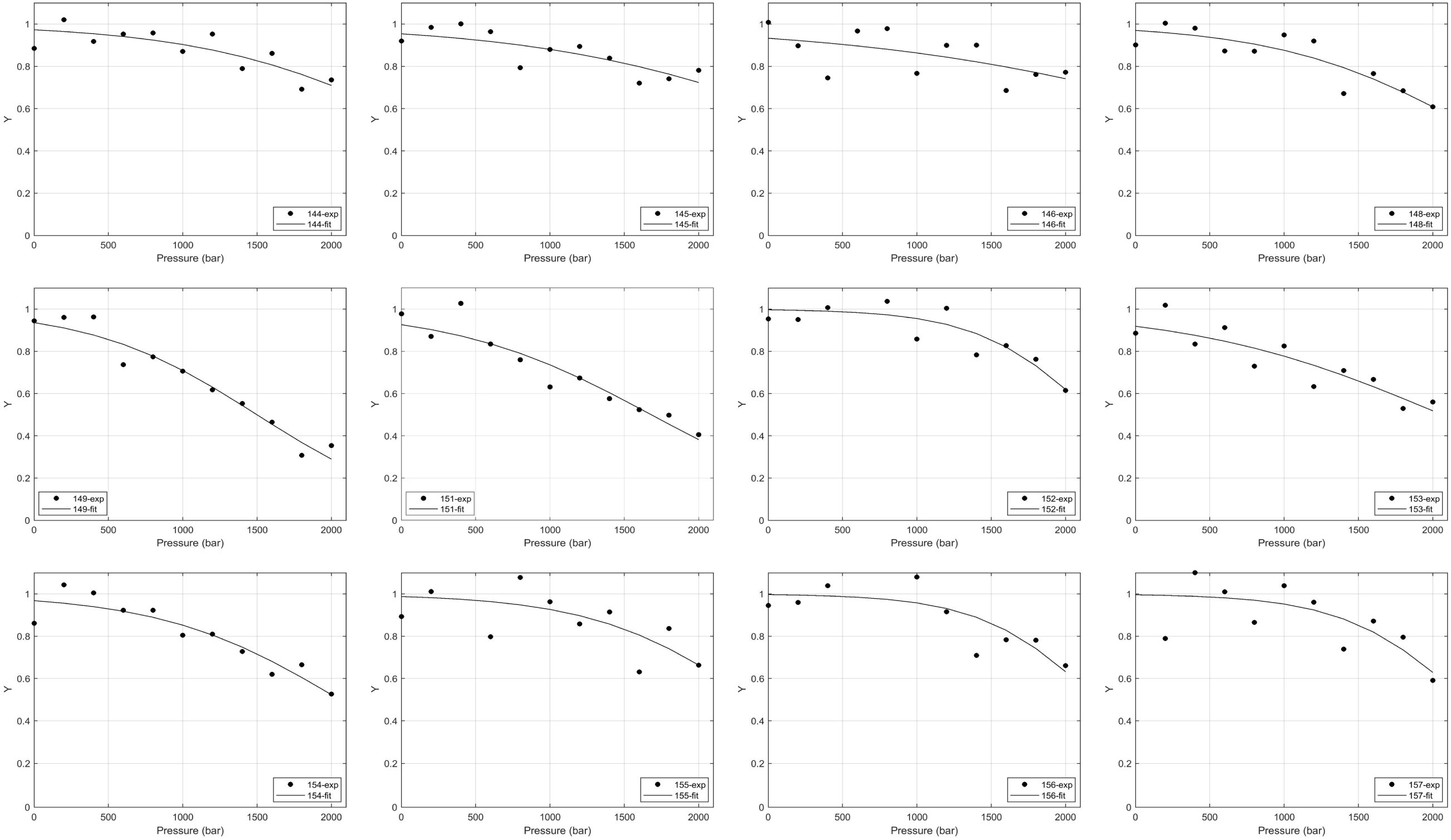
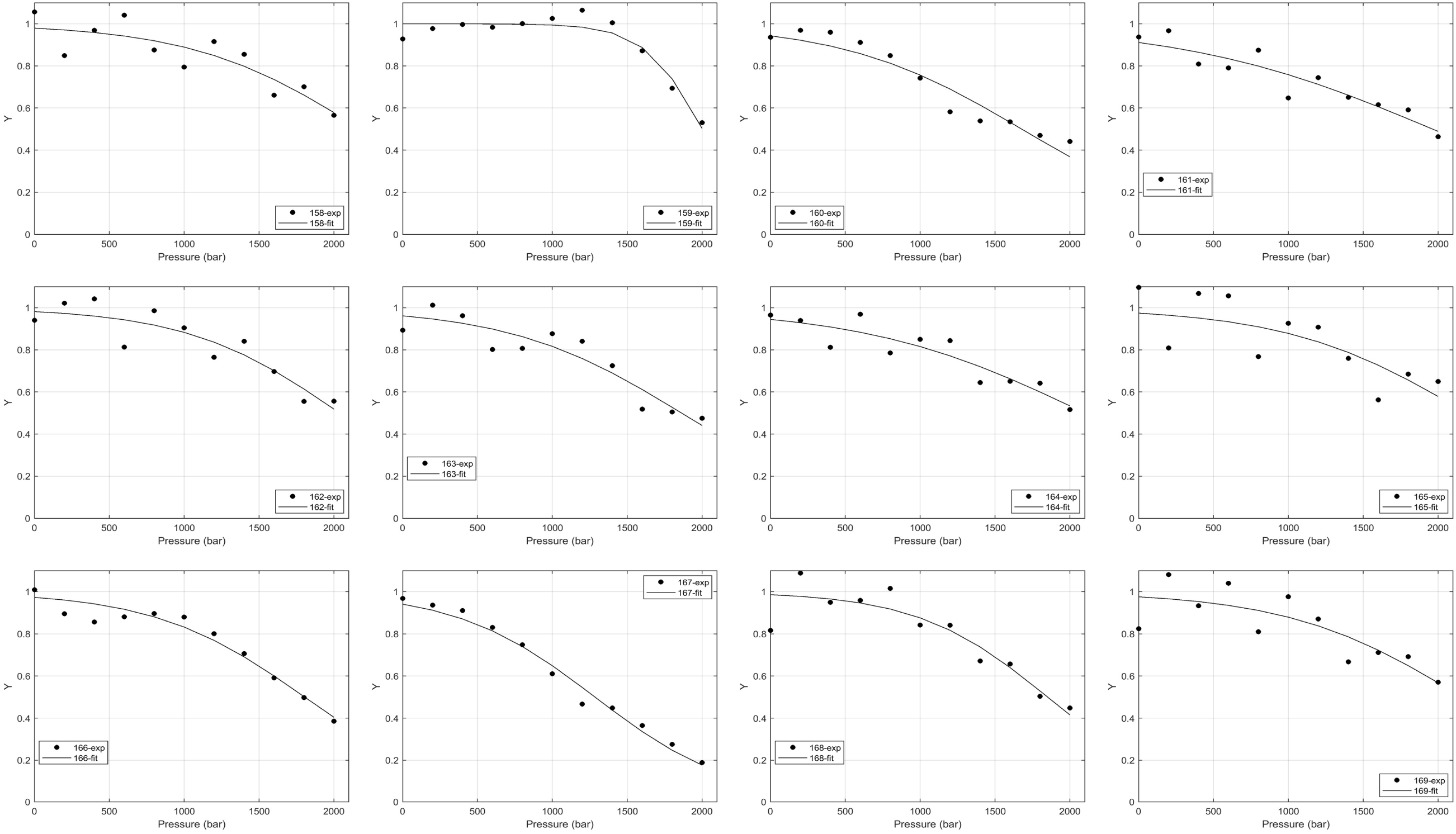
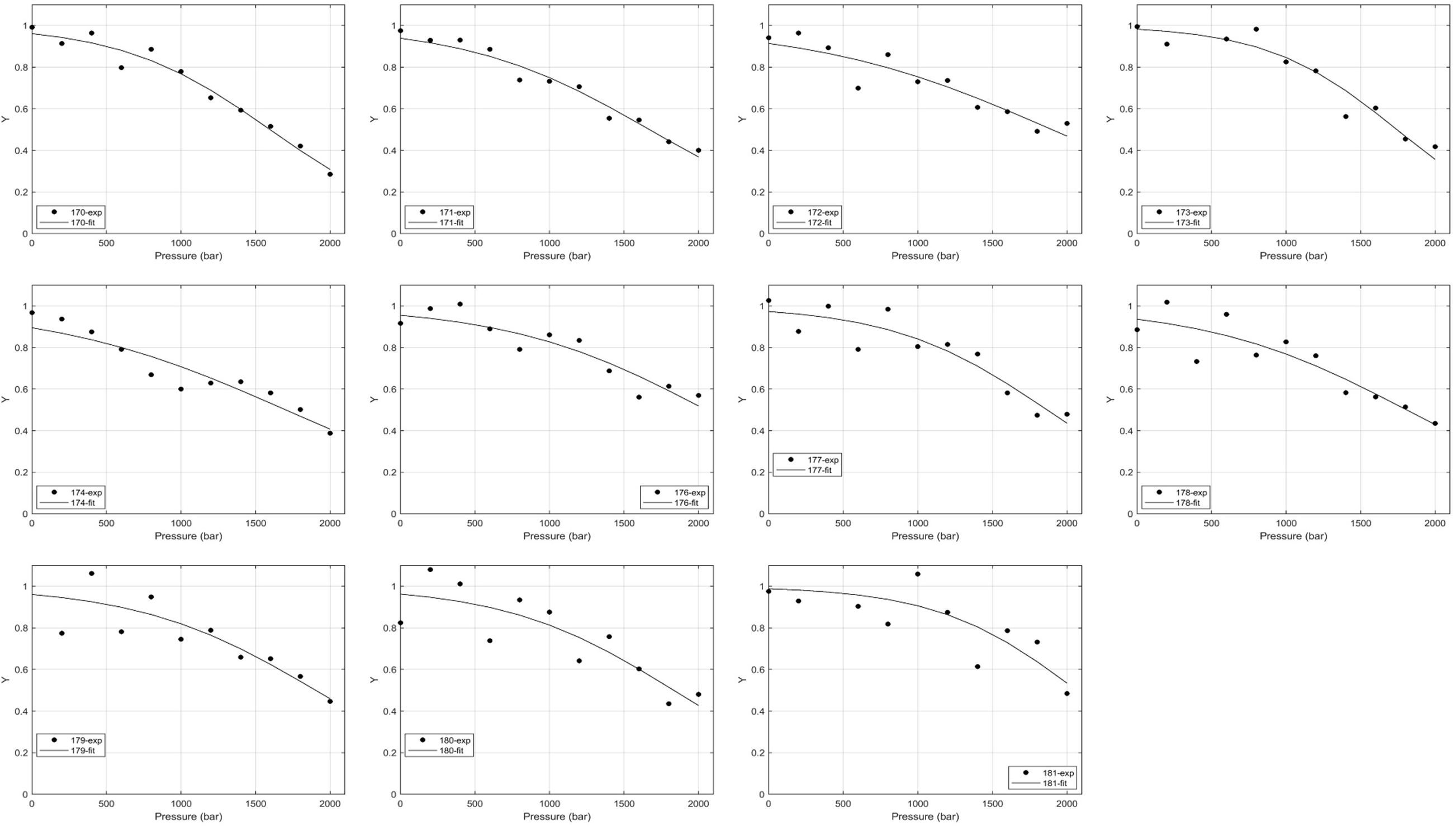
Individual transition curves for all quantified residues. Black circles are normalized data points. Black lines are the corresponding fits. Raw data were fit and then both data and fits were normalized for comparison.

**Figure S5.**
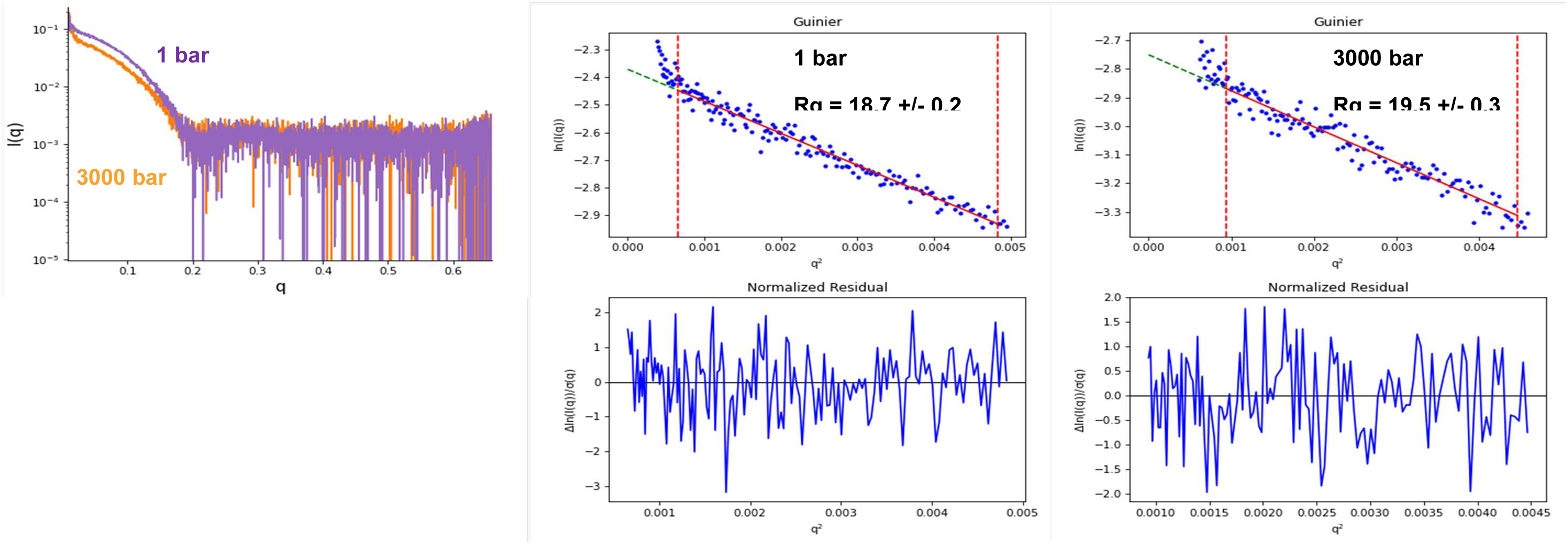
Pressure-dependent SAXS profiles for Arf1 I42S. Left) Buffer corrected scattering intensity profiles for 1 bar and 3000 bar as indicated. Middle) Guinier plots for Arf1 I42S scattering profile at 1 bar. Lower panel shows residuals of the Guinier fits. The Rg value is consistent with the size and shape of folded Arf proteins. Right) Arf1 I42S scattering profile at 3000 bar. Lower panel shows residuals of the Guinier fits. The deviation at low q is due to imperfect buffer subtraction.

